# Essential role of PLD2 in hypoxia-induced stemness and therapy resistance in ovarian tumors

**DOI:** 10.1101/2023.07.19.549738

**Authors:** Sandra Muñoz-Galván, Eva M. Verdugo-Sivianes, José M. Santos-Pereira, Purificación Estevez-García, Amancio Carnero

## Abstract

Hypoxia in solid tumors is a source of chemoresistance that determines poor patient prognosis and relies on the presence of cancer stem cells (CSCs). Here we use ovarian cancer (OC) as a model and a combination of 2D and 3D cell cultures, xenograft models, patient samples, transcriptional databases, iPSCs and ATAC-seq, to address the mechanisms leading to hypoxia-induced CSC generation and chemoresistance. We show that hypoxia activates the expression of the PLD2 gene encoding phospholipase D2. PLD2 overexpression leads to increased CSC-like features, similar to hypoxia, while PLD2 depletion in hypoxia partially suppresses these effects, indicating a role of PLD2 in hypoxia-induced CSC generation in OC. Finally, PLD2 overexpression provokes chemoresistance that is suppressed by combination treatment with PLD2 inhibition. Altogether, our work highlights the HIF-1D-PLD2 axis in hypoxia-induced CSC generation and chemoresistance in OC and proposes an alternative treatment for patients with high PLD2 expression.

**Statement of Significance:** Hypoxia in solid tumors is a major source of chemoresistance and cancer stem cells. We show that hypoxia-induced stemness is mediated by phospholipase D2 in ovarian tumors, generating therapy resistance that is overcome by phospholipase D inhibition. Therefore, we propose an alternative treatment for patients with high *PLD2* expression.

## INTRODUCTION

Hypoxia is a common feature of the tumor microenvironment in solid tumors ^1^. Hypoxia is generated by insufficient oxygen diffusion towards parts of the tumor that are not irrigated, although the highly irregular tumor microvasculature may also generate hypoxic regions. Cancer cells under hypoxic conditions undergo a series of transcriptional changes that are induced by hypoxia-inducible factors (HIF), HIF-1, –2 and –3, with HIF-1 being the best known. HIFs are heterodimeric transcription factors containing bHLH-PAS domains and consist of an α subunit regulated by oxygen levels and a constitutively expressed β subunit (also called ARNT). The HIFa-ARNT dimers proteins bind DNA at specific sequences known as hypoxia-response elements (HREs) to promote targeted gene expression ^2^. In normoxia, HIF-α subunits are hydroxylated at specific proline and asparagine residues by prolyl hydroxylase 2 (PHD2) and are recognized and targeted for degradation by the von Hippel‒Lindau (VHL) tumor suppressor. However, hypoxia leads to the inhibition of such hydroxylation and subsequent HIF-α accumulation, heterodimerization with ARNT and transcriptional activation of target genes ^3^. The multiple effects of hypoxia on the biology of tumors include preventing apoptosis and promoting proliferation and autophagy, inducing metabolic alterations, and promoting angiogenesis, the epithelial-to-mesenchymal (EMT) transition, invasion and metastasis ^4, 5^. Indeed, HIF-1α is overexpressed in multiple tumor types and is associated with a poor prognosis in patients and therapy resistance ^6–13^.

A major cause of chemoresistance is the persistence of a cancer cell subpopulation known as cancer stem cells (CSCs) or tumor-initiating cells. CSCs are able to recapitulate a new tumor since they possess self-renewal and pluripotency properties similar to those of normal stem cells ^14, 15^. CSCs are resistant to common antitumor therapies, which may indeed cause their enrichment, leading to chemoresistance and relapse ^16, 17^. Several studies indicate that HIF-1 is required for maintaining CSCs and that its activation in hypoxia leads to the increased expression of stem marker genes in multiple cancer types ^18, 19^. For instance, in glioma CSCs, stemness is reduced upon either *HIF1α* or *HIF2α* silencing in both normoxic and hypoxic environments ^20^. In a breast cancer model, the conditional deletion of *Hif1α* in mammary epithelial cells led to a reduced CSC population and function, causing less tumor growth and metastasis ^21^. HIF factors are also important for the maintenance of CSCs in haematological malignancies, such as lymphoma and acute myeloid leukaemia ^22^. Therefore, clearly, hypoxia promotes CSCs and leads to chemoresistance through HIF factors; however, knowledge regarding HIF targets that may be responsible for CSC activity is limited.

Ovarian cancer (OC) is the most lethal gynaecological malignancy ^23^ mainly due to its nonspecific clinical manifestations, which lead to a late diagnosis and high chemoresistance ^24^. The standard treatment for OC consists of cytoreductive surgery, followed by chemotherapy with platinum-based compounds alone or in combination with other anticancer drugs ^25^. However, due to the high rates of relapse (80% in advanced stages and 20% in early stages), OC is incurable in most cases. Similar to other solid tumors, hypoxia is a key microenvironment modulator in OC, affecting not only the primary tumor but also the main location of OC metastases, which are abdominal ascites ^8, 26^. OC cell lines subjected to hypoxic conditions behave more aggressively and overexpress stem cell markers ^27^, and upregulated sirtuin 1 expression has been linked to ovarian CSCs through HIF-1α ^28^. Therapeutic resistance has also been reported in OC under hypoxic conditions, and HIF-1α may upregulate genes encoding the transporter ABCG2 to increase drug efflux ^29, 30^ and induce alterations in metabolism that compromise sensitivity to cisplatin ^31^. However, the mechanisms responsible for the generation of the CSC phenotype under hypoxic conditions in OC remain elusive.

Phospholipases D (PLD) are enzymes that hydrolyse phosphatidylcholine to produce phosphatidic acid (PA) and choline, playing multiple cellular roles, including cell proliferation, protein trafficking, membrane remodelling and cytoskeletal dynamics ^32^. There are two main paralogs as follows: PLD1 and PLD2 ^33, 34^. *PLD2* is overexpressed in multiple cancer types, being activated by oncogenes, such as Ras, Rho, Src, Raf and Erk, and is related to increased proliferation, adhesion, invasion and metastasis in a mechanism that involves its enzymatic product PA, interactions with proteins that result in increased actin polymerization and activation of mammalian target of rapamycin (mTOR) ^35–40^. We previously showed that *PLD2* is overexpressed in colorectal cancer patients and that its effect on tumorigenesis is achieved by communication with the tumor microenvironment in a feedback loop ^41^. PLD2 and PA secreted by cancer cells induce senescence in stromal fibroblasts, which, in turn, produce a battery of cytokines and growth factors that enhance the stemness of cancer cells. Recent evidence showed that *PLD2* expression is induced by HIF-1α in colon cancer cells ^42^, and conversely, PLD2 is able to activate HIF-1α in glioma and renal cells through PA ^43, 44^. Additionally, the deletion of *PLD2* in endothelial cells reduces the hypoxic response and angiogenesis ^45^. These data suggest the existence of a positive feedback loop in which PLD2 and HIF-1α potentiate each other, although this could be context dependent since PLD2 has a negative impact on HIF-1α in other cell types ^46^. In OC, *PLD2* expression in high-grade serous carcinoma is higher in effusions than in solid tumors or metastases ^47^, suggesting its role in promoting invasiveness. However, there is a lack of functional evidence showing a role of PLD2 in tumorigenesis and stemness in OC.

In this work, we show that hypoxia induces an increase in *PLD2* expression in a HIF-1α-dependent manner. The overexpression of *PLD2* in OC cells and xenograft models leads to increased CSC features, while PLD2 depletion has the opposite effects, indicating a major role of PLD2 in the generation of ovarian CSCs. This effect was also observed in cell reprogramming experiments with iPSCs. Importantly, the increase in CSC-like features induced by hypoxia partially relies on *PLD2* expression. PLD2 is also required for chromatin accessibility and stemness gene expression induced by hypoxia. The expression of *PLD2* is increased in OC tumors, thereby leading to the rewiring of transcriptional programmes related to stem cell proliferation and maintenance and response to hypoxia. Finally, we show both *in vitro* and *in vivo* that *PLD2* overexpression causes chemoresistance to platinum-based drugs, whereas *PLD2* depletion suppresses such resistance, indicating that *PLD2* overexpression is a prognostic factor for tumor aggressiveness and chemoresistance in OC. We also propose an alternative treatment for patients overexpressing *PLD2,* consisting of the combination of cisplatin and the pharmacological inhibition of PLD2, which is able to suppress therapy resistance. Altogether, our data show that the HIF-1α-PLD2 axis is an important player in OC biology that could be used as a therapeutic target to overcome therapy resistance.

## RESULTS

### Hypoxia induces a CSC-like phenotype in OC cells

The ability of hypoxia to induce a CSC-like phenotype in OC cells has been previously observed ^27^. We first aimed to validate these results in our OC cell lines SKOV3, OVCAR8 and ES-2 and found that hypoxic conditions of 3% O_2_ led to significant increases in the number of tumorspheres generated by growing the cells under low-attachment conditions and in the percentage of holoclones, both of which were used as a proxy for CSCs (Supplementary Figure S1A-B). Next, we analysed the expression of stem cell markers in OC cell lines grown under hypoxic conditions and detected an increase in the mRNA levels of *NANOG*, *CD44*, *SOX2* and *EPCAM* and the percentage of cells containing the surface CSC marker CD133 (Supplementary Figure S1C-D). These results confirm that hypoxia induces a CSC-like phenotype in OC cells.

### Hypoxia leads to the overexpression of *PLD2* through HIF-1α in OC cells

We previously showed that PLD2 is involved in communication with the tumor microenvironment in colorectal cancer, and its overexpression leads to increased tumor stemness ^41^. In addition, *PLD2* expression is increased under hypoxic conditions in colon cancer cells ^42^. Therefore, we wondered whether hypoxia-induced stemness of OC cells could be mediated by PLD2. *PLD2* expression levels in cell lines from gynecological cancers are heterogeneous, independent on *tp53* mutation status and associated to particular histotypes, being higher in germ cell carcinomas, medium in serous and endometroid carcinomas and lower in clear cell carcinomas (Supplementary Figure S2). We selected SKOV3, OVCAR8 and ES-2 OC cell lines for our studies and first analysed the expression levels of *PLD2* in OC cells under hypoxic conditions. We found that *PLD2* expression was similar among the three cell lines and that oxygen levels of 3% led to a 2-fold increase in the expression of *PLD2* in all of them, while the well-known hypoxia target genes *LDHA* and *VEGFA* showed a similar increase, validating the results (Figure 1A). The hypoxia-induced increase in PLD2 expression was also observed at the protein level by performing immunofluorescence in the three OC cell lines (Figure 1B). In addition, we further demonstrated these findings by using the HIF-hydroxylase inhibitor DMOG, which increases HIF levels, generating a hypoxia-like phenotype under normoxic conditions (Supplementary Figure S3A-B). Thus, we treated OC cells with DMOG and found that this treatment resulted in a similar increase in hypoxia marker gene expression as that induced by hypoxia, also leading to the observed increase in *PLD2* expression at both the mRNA and protein levels (Figure 1A-B).

**Figure 1.**
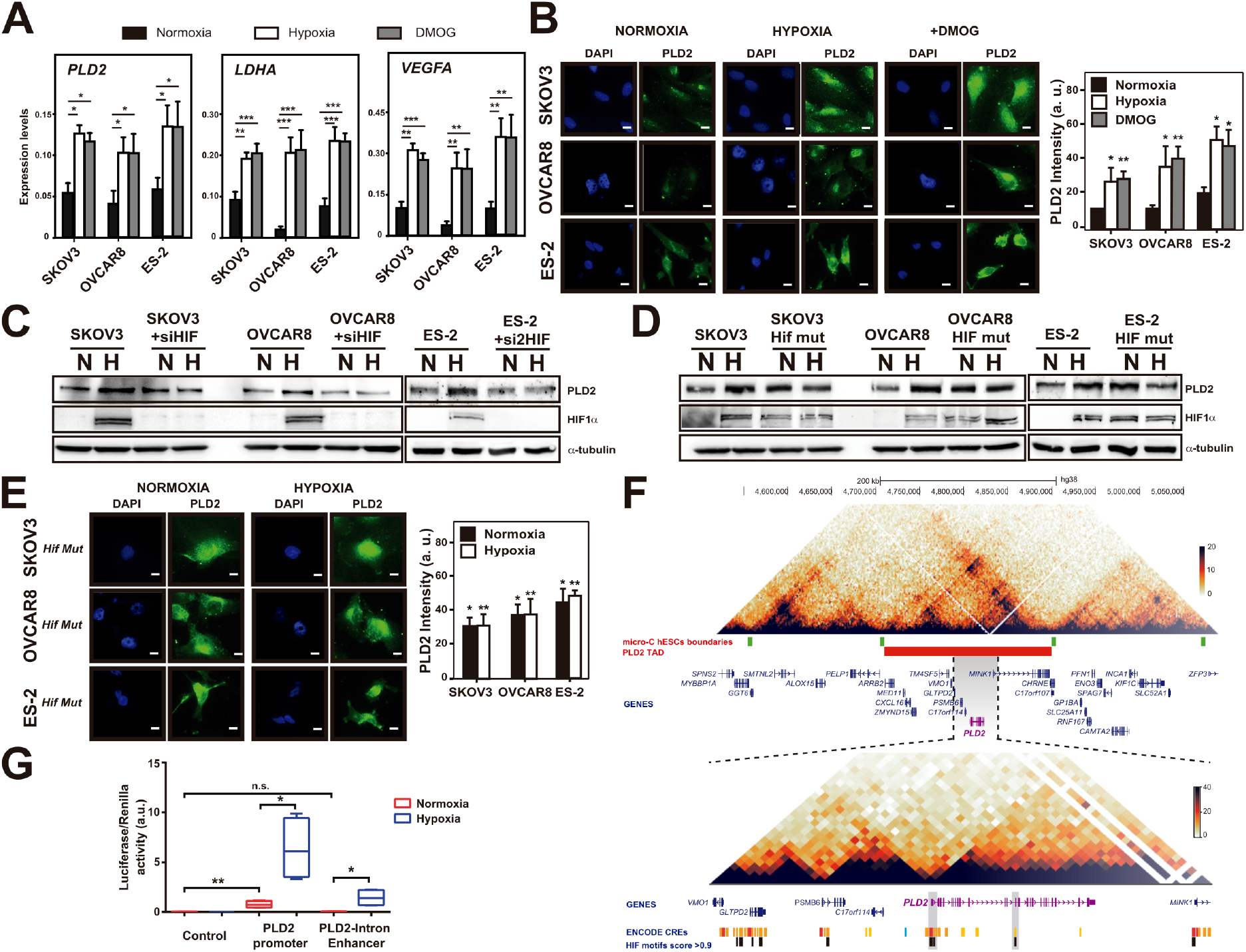
Hypoxia leads to the overexpression of *PLD2* through HIF-1α in OC cells. (**A**) *PLD2*, *LDHA* and *VEGFA* expression levels in SKOV3, OVCAR8 and ES-2 ovarian cancer cell lines under normoxia (black) and hypoxia (white) or in the presence of the HIF hydroxylase inhibitor DMOG (grey), measured by RT-qPCR. The mRNA expression was calculated as 2^-ΔCt^ relative to the *ACTB* gene. **(B)** Representative images and quantification of PLD2 protein levels by immunofluorescence in SKOV3, OVCAR8 and ES-2 cells carrying empty vector (Ev) under normoxia (black) and hypoxia (white) or in the presence of the HIF-hydroxylase inhibitor DMOG (grey). Scale bars: 10 μm. **(C)** Western blot showing PLD2, HIF-1α, and alpha-tubulin protein levels in SKOV3 or OVCAR8 ovarian cancer cells in the presence or absence of a small interfering RNA (siRNA) of HIF-1α. **(D)** Western blot showing PLD2, HIF-1α, and alpha-tubulin protein levels in SKOV3 and OVCAR8 cells expressing a *hif1a* mutant (HIF mut) or Ev. **(E)** Representative images and quantification of PLD2 protein levels by immunofluorescence in SKOV3, OVCAR8 and ES-2 cells expressing a *hif1a* mutant (Hif Mut) under normoxia (black) and hypoxia (white). Scale bars: 10 μm. **(F)** Top, heatmaps showing the microC signal in hESCs in a 0.5-Mb region of chromosome 17, microC hESC boundaries (green), TAD containing the *PLD2* gene (red) and genes (blue). Bottom, zoom within the TAD containing the *PLD2* gene showing ENCODE cis-regulatory elements (CREs, yellow to red) and those containing HIF1 motifs with relative scores higher than 0.9 (black). **(G)** Luciferase activity assay of the *PLD2* promoter and putative enhancer in HEK293 cells under hypoxic or normoxic conditions. A minimum of three biological were performed per each experiment. The data were compared using Student’s t tests. For A, B and G, asterisks indicate statistical significance with respect to normoxia. For E, asterisks indicate statistical significance with respect to normoxia in panel B. *p < 0.05; **p < 0.01; ***p < 0.001.

Next, we wondered whether the hypoxia-induced upregulation of PLD2 was indeed mediated by HIF-1α. Therefore, we depleted *HIF1A* using a small interfering RNA (siRNA) in the three OC cell lines and found that it suppressed the increase in the PLD2 protein levels induced by hypoxia (Figure 1C). Additionally, we generated *hif1a* mutant OC cell lines by transfecting cells with a mutant *hif1a* allele ^48^ that is unable to be hydroxylated and, therefore, is constitutively active even under normoxic conditions. Interestingly, we observed that the PLD2 levels in the *hif1a* mutant cells were as high as those induced by hypoxia on a *HIF1α* wild-type background in both normoxia and hypoxia, confirming that HIF-1α activates PLD2 expression (Figure 1D-E). Altogether, these data indicate that hypoxia induces the upregulation of *PLD2* expression in OC cells that is mediated by HIF-1α.

### HIF-1α activates *PLD2* transcription through HREs at promoter and hypoxia-specific enhancer regions

We aimed to understand how HIF-1α promotes *PLD2* expression. HIF-1α is considered the master transcriptional regulator of the cellular response to hypoxia. HIF-1α forms a heterodimer with ARNT that binds HREs to control the expression of hypoxia-response genes^2^. To address the possibility that HIF-1α could regulate *PLD2* expression at the transcriptional level, we first searched for possible cis-regulatory elements (CREs) near the *PLD2* gene that may contain HREs. According to public 3D chromatin conformation experiments (micro-C) in human embryonic stem cells, the *PLD2* gene is located within a topologically associating domain (TAD) of 190 kb with a high interaction frequency at the 3D level and is relatively isolated from the neighbouring regions (Figure 1F). To identify CREs that may regulate *PLD2* expression, we focused on a smaller region of 50 kb surrounding the *PLD2* gene with a higher interaction frequency with the *PLD2* promoter, which we called the *PLD2* regulatory region (Figure 1G). We scanned the CREs annotated by ENCODE within the *PLD2* regulatory region for the presence of the DNA binding motif of HIF1A with a high score (>90% relative score). We found 15 of 35 CREs fulfilling this condition, some of which corresponded to gene promoters and others to enhancers, including the *PLD2* promoter and an enhancer in *PLD2* intron 12 (Figure 1G). These CREs with high-score HIF1A motifs represent putative HREs.

To assess the regulatory activity of the two CREs in the *PLD2* gene (promoter and putative enhancer) containing the HIF1A motif, we cloned both genomic regions in promoter and enhancer reporter vectors while controlling the expression of the luciferase gene. We transfected OC cells with these vectors and measured the luciferase activity under normoxia and hypoxia. We found that the *PLD2* promoter was able to activate reporter expression in normoxia and that hypoxia led to a significant increase in luciferase activity (8-fold) (Figure 1H), suggesting that the *PLD2* promoter responds to hypoxic conditions by increasing *PLD2* transcription. However, the enhancer contained within the *PLD2* intron was unable to activate reporter expression in normoxia but led to a 22-fold increase in its expression in hypoxia (Figure 1H), indicating that this enhancer acts as an HRE in OC cells.

### *PLD2* promotes tumorigenesis and CSC-like features in OC cells

Since hypoxia has been linked to increased CSC features in several cancer types ^18, 19^, we wondered whether the *PLD2* increase in expression led to an increase in the CSC population in OC cells. Therefore, we first established OC cell lines expressing ectopic *PLD2* cDNA or depleted *PLD2* using a short hairpin RNA (shRNA). The expression of *PLD2* under these conditions was assessed at the mRNA and protein levels (Figure 2A-B and Supplementary Figure S3C). We observed that the overexpression of *PLD2* in OC cells led to a significant increase in the number of clones generated by the three cell lines, while a significant decrease was detected in the OVCAR8 cells upon the *PLD2* depletion (Figure 2C), suggesting that PLD2 promotes tumor growth. To address this question, we analysed the growth of these cell lines and found that the enhanced *PLD2* expression led to a significant increase in proliferation, while the *PLD2* depletion generated the opposite effect with statistical significance in all cell lines (Figure 2D). This effect was further confirmed *in vivo* by generating xenograft models of OC cells overexpressing or depleted of *PLD2*, showing an increase or decrease in the tumor volume, respectively, 50 days after transplantation (Figure 2E). Altogether, these results indicate that *PLD2* expression promotes tumorigenesis.

**Figure 2.**
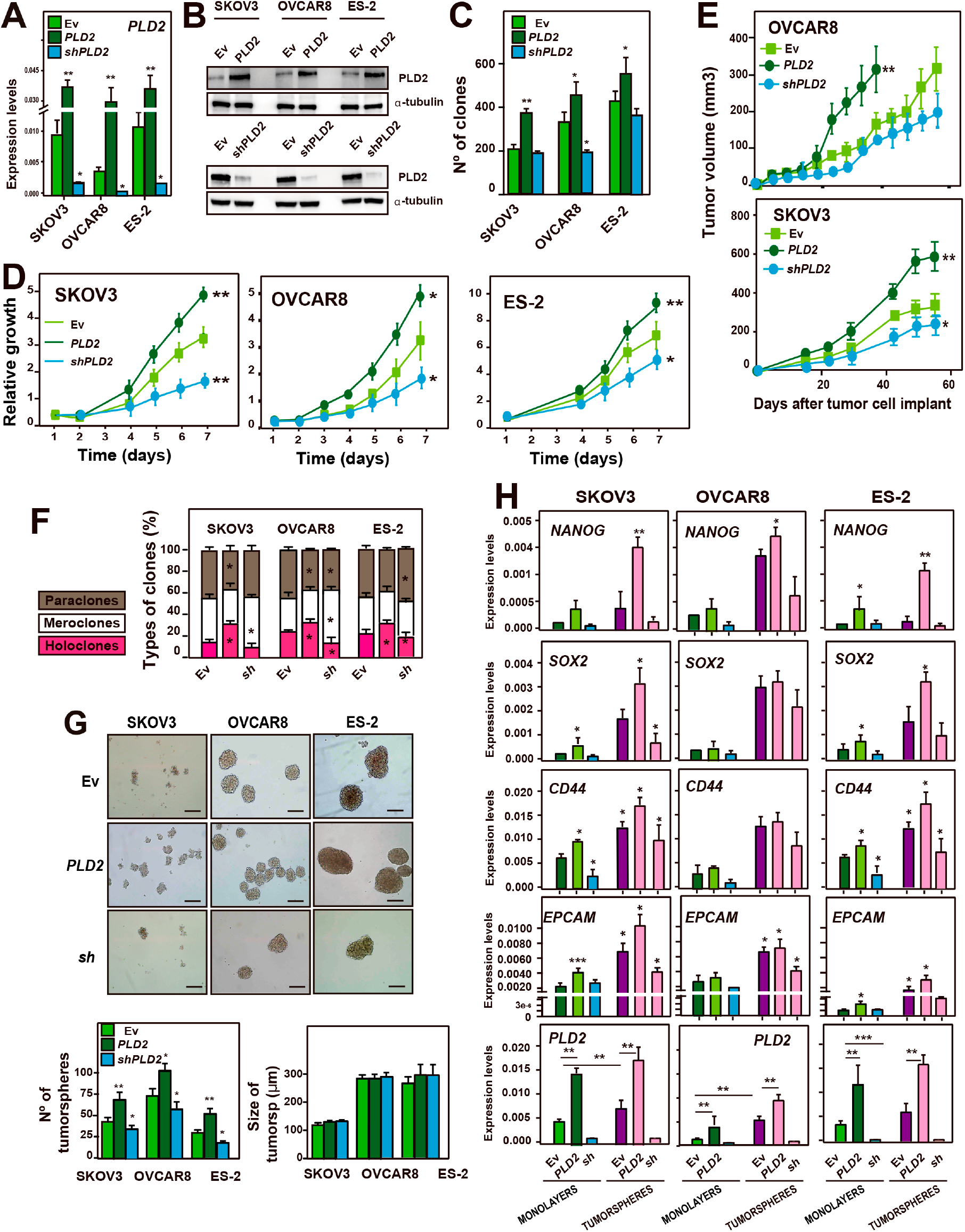
*PLD2* promotes tumorigenesis and CSC-like features in ovarian cancer. (**A**) Analysis of the expression of *PLD2* by RT‒qPCR in SKOV3, OVCAR8 and ES-2 cells carrying an empty vector (Ev) expressing *PLD2* or *shPLD2*. The mRNA expression was calculated as 2^-ΔCt^ relative to the *ACTB* gene. **(B)** Western blot showing the protein levels of PLD2 in cells carrying Ev and expressing *PLD2* or *shPLD2*. **(C)** Quantification of the number of colonies formed by SKOV3, OVCAR8 and ES-2 cells carrying Ev and expressing *PLD2* or *shPLD2*. **(D)** Growth curve of SKOV3, OVCAR8 and ES-2 cells carrying Ev (light green) or expressing *PLD2* (dark green) or *shPLD2* (blue) represented as the accumulation of the doubling times. **(E)** Tumor growth in xenografts from SKOV3 and OVCAR8 cells carrying Ev, overexpressing *PLD2* or *shPLD2* were injected into female athymic nude mice. Cohorts of 5 mice each were used. **(F)** Percentage of paraclones, meroclones and holoclones formed by SKOV3, OVCAR8 and ES-2 cells carrying Ev expressing *PLD2* or *shPLD2*. At least 200 individual clones were analysed. The averages and SDs of three independent experiments are shown. **(G)** Top, Representative images of tumorspheres formed by SKOV3, OVCAR8 and ES-2 cells carrying Ev expressing *PLD2* or *shPLD2*. Bottom, quantification of the number and size of tumorspheres. Scale bars: 250 μm. **(H)** Analysis of the expression by RT‒qPCR of stemness-associated genes and *PLD2* in total cell extracts and tumorspheres from SKOV3, OVCAR8 and ES-2 cells carrying Ev and expressing *PLD2* or *shPLD2*. The mRNA expression was calculated as 2^-ΔCt^ relative to the *ACTB* gene. The average and SD of at least three independent experiments are shown. The data were analysed using Student’s t test. Asterisks indicate statistical significance with respect to Ev carrying cells. *, P < 0.05; **, P < 0.01; ***, P < 0.001.

Next, we wondered whether the *PLD2* expression levels were related to the formation of ovarian CSCs. First, we analysed the formation of different types of colonies, including holoclones, meroclones and paraclones, which are considered stem cells, transit-amplifying cells and differentiated cells, respectively ^49^. We found a significant increase in the percentage of holoclones and a significant decrease in the percentage of paraclones in the three cell lines overexpressing *PLD2* (Figure 2F). A significant decrease in holoclone formation was also observed in the three OC cell lines upon the *PLD2* depletion. Furthermore, we measured the formation of tumorspheres under low attachment conditions, which represent tumor initiating cells, in OC cells overexpressing or depleted of *PLD2*. We found that *PLD2* overexpression led to a significant increase in the number of tumorspheres, while *PLD2* depletion generated the opposite effect (Figure 2G), although we did not observe changes in the size of such tumorspheres. These data indicate that *PLD2* expression promotes the formation of ovarian CSCs.

Finally, we analysed the expression of pluripotency and CSC marker genes in our OC cell lines overexpressing or depleted of *PLD2*. We found that the expression of *SOX2*, *CD44* and *EPCAM* was significantly increased in the ES-2 and SKOV3 cells overexpressing *PLD2*, while *NANOG* was only increased in ES-2 cells (Figure 2H). In OVCAR8 cells, the expression of these genes was increased in the same trend, although in a nonsignificant manner. Then, we measured the levels of expression of these pluripotency genes in the tumorspheres extracts. First, we found that *PLD2* was highly expressed in the tumorspheres compared with that in the total extracts transfected with only the empty vector (Figure 2H), suggesting that CSCs indeed have higher expression levels of *PLD2*. The expression of stemness genes was also increased in the tumorspheres compared with that in the total extracts, as expected, while they were further upregulated in most cases upon the *PLD2* overexpression or downregulation under *PLD2* depletion (Figure 2H). Altogether, these data indicate that *PLD2*, whose expression is induced by hypoxia, is an important oncogene in OC and that its overexpression leads to increased tumor stemness.

### The hypoxia-induced stemness of OC cells partially depends on *PLD2* expression

Thus far, we showed that both hypoxia and *PLD2* expression led to an increase in CSCs in OC cells (Figure 2, Supplementary Figure S1) and that *PLD2* expression was increased under hypoxic conditions in a HIF-1α-dependent manner (Figure 1). Thus, we wondered whether both phenomena were connected and whether the hypoxia-induced increase in CSC-like features was dependent on *PLD2* expression. To address this question, we first analysed the formation of tumorspheres in normoxia and hypoxia following a reduction in *PLD2* levels. We observed that either *PLD2* overexpression or hypoxia led to a similar increase in the formation of tumorspheres, with only a slightly higher nonsignificant increase when both conditions were combined (Figure 3A). However, the *PLD2* decrease by shRNA caused a partial suppression of the increase in tumorsphere formation under hypoxic conditions in SKOV3 and OVCAR8 cells, which was rescued by overexpressing back *PLD2* in shRNA-transfected cells (Figure 3A and Supplementary Figure S4A), suggesting that PLD2 is partially responsible for the hypoxia-induced stemness. Then, we measured the formation of holoclones, meroclones and paraclones under hypoxic conditions upon *PLD2* overexpression or depletion. We found that the increase in the percentage of holoclones induced by hypoxia was further enhanced by the *PLD2* overexpression, while it was suppressed by the *PLD2* depletion and rescued back by expressing *PLD2* after its depletion (Supplementary Figure S4B).

**Figure 3.**
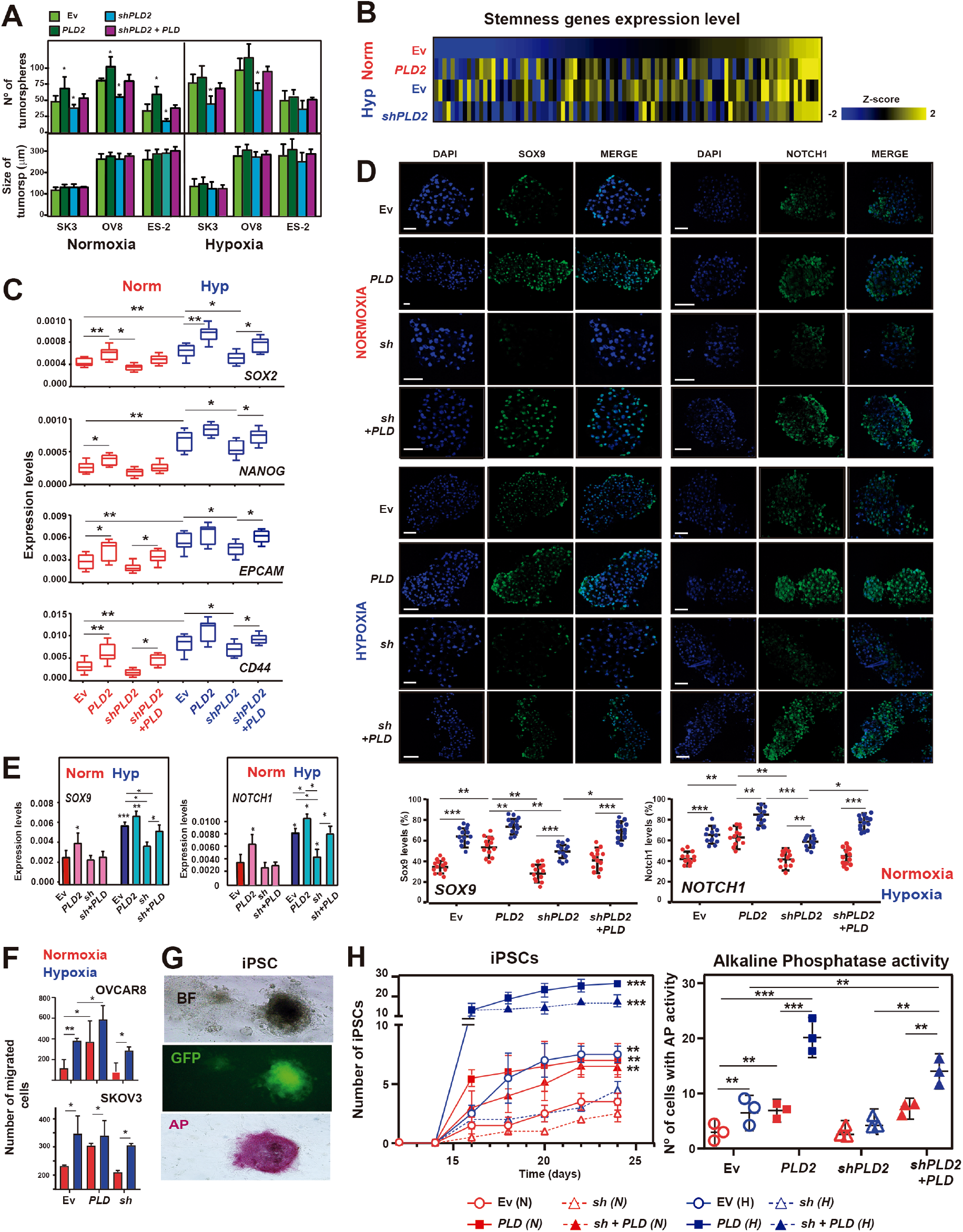
The hypoxia-induced stemness of OC cells partially depends on PLD2 expression. (**A**) Quantification of the number and size of tumorspheres formed by SKOV3, OVCAR8 and ES-2 cells carrying Ev and expressing *PLD2*, *shPLD2* or both. **(B)** Heatmaps showing the z scores of the expression of stemness-associated genes obtained from TaqMan arrays. Genes are sorted according to decreasing z scores in Ev-carrying cells under normoxia. **(C)** Expression levels of *SOX2, NANOG, CD44* and *EPCAM* stemness-associated genes in SKOV3, OVCAR8 and ES-2 cells carrying Ev and expressing *PLD2*, *shPLD2* or both under normoxic or hypoxic conditions. The mRNA expression was calculated as 2^-ΔCt^ relative to the *ACTB* gene. **(D)** Top, determination of the Sox9 and Notch1 protein levels by immunofluorescence in tumorspheres formed by OC cell lines carrying Ev and expressing *PLD2*, *shPLD2* or both. Bottom, quantification of the percentage of cells with Sox9 and Notch1 expression in tumorspheres formed by ovarian cancer cell lines carrying Ev and expressing *PLD2* or *shPLD2*. Scale bars: 100 μm. **(E)** Expression levels of *SOX9* and *NOTCH1* stemness-associated genes in tumorspheres formed by ovarian cancer cell lines carrying Ev and expressing *PLD2*, *shPLD2* or both. The mRNA expression was calculated as 2^-ΔCt^ relative to the *ACTB* gene. **(F)** Quantification of Boyden chamber migration assays in SKOV3 and OVCAR8 cells carrying Ev and expressing *PLD2* or *shPLD2* under normoxic or hypoxic conditions. **(G)** Representative images of induced pluripotent stem cells (iPSCs) by reprogramming mouse embryonic fibroblasts (MEFs) with OSKM genes (Yamanaka factors Oct3/4, Sox2, Klf4 and cMyc) and Nanog reporter retroviruses. Top, bright field microscopy of iPSCs. Medium, immunofluorescence image showing GFP in cells in which *Nanog* is active. Bottom, Alkaline phosphatase activity. **(H)** Left, quantification of iPSCs generated from MEFs infected with OSKM genes and an additional lentiviral vector that expresses GFP in cells in which the Nanog promoter/enhancer is active and the corresponding plasmid overexpressing *PLD2*, *shPLD2*, both or carrying Ev under normoxia or hypoxia. Right, quantification of cells with alkaline phosphatase activity at the end of the iPSC generation experiment. A minimum of three biological were performed per each experiment. The data were analysed using Student’s t test. Asterisks indicate statistical significance with respect to Ev carrying cells, unless indicated by horizontal lines. *, P < 0.05; **, P < 0.01; ***, P < 0.001.

Next, we analysed the expression levels of stemness genes by RT‒qPCR using custom TaqMan Array plates containing probes against a selection of these genes in ovarian cancer cells. We observed that either hypoxia or *PLD2* overexpression in normoxia led to the increased expression of many stemness genes, while the *PLD2* depletion largely suppressed this increase (Figure 3B). We confirmed this result by RT‒qPCR of individual representative genes, including *SOX2*, *NANOG*, *CD44* and *EPCAM*, showing that either hypoxia or *PLD2* overexpression in normoxia lead to increased expression of stemness genes, while combination of both conditions further increased their expression (Figure 3C). *PLD2* depletion partially suppressed the hypoxia-induced enhancement of expression, with a lower non-statistically significant effect in normoxia, and rescue experiments confirmed the specificity of *PLD2* depletion (Figure 3C). Interestingly, hierarchical clustering of the four analysed conditions showed that the samples corresponding to EV hypoxic cells and *PLD2*-overexpressing cells clustered together, while EV normoxic cells and *PLD2*-depleted cells clustered separately (Supplementary Figure S5A).

Then, we measured the protein levels of the pluripotency factors Sox2, Sox17, Sox9 and Notch1 (found to correlate with PLD2 in tumors, see below Figure 4E) in tumorspheres by immunofluorescence to determine whether PLD2 could influence their expression in CSCs. We validated PLD2 protein levels in tumorspheres (Supplementary Figure S5B) and found that either hypoxia or *PLD2* overexpression led to an increase in the levels of Sox2, Sox9 and Notch1, while only hypoxia led to an increase in the Sox17 protein levels (Figure 3D and Supplementary Figure S5C). In addition, the *PLD2* depletion led to a partial suppression of the hypoxia-induced expression of Sox2, Sox9 and Notch1 that was rescued by expressing back *PLD2* in these cells (Figure 3D and Supplementary Figure S5C), suggesting that PLD2 plays a role in the generation of CSCs in hypoxia through these genes. These observations were confirmed at the mRNA level by RT‒qPCR (Figure 3E; Supplementary Figure S5D). Altogether, these data indicate that PLD2 plays a major role in the induction of the CSC phenotype in hypoxia, promoting the expression of specific stem-related genes, such as *SOX2*, *SOX9* or *NOTCH1*.

**Figure 4.**
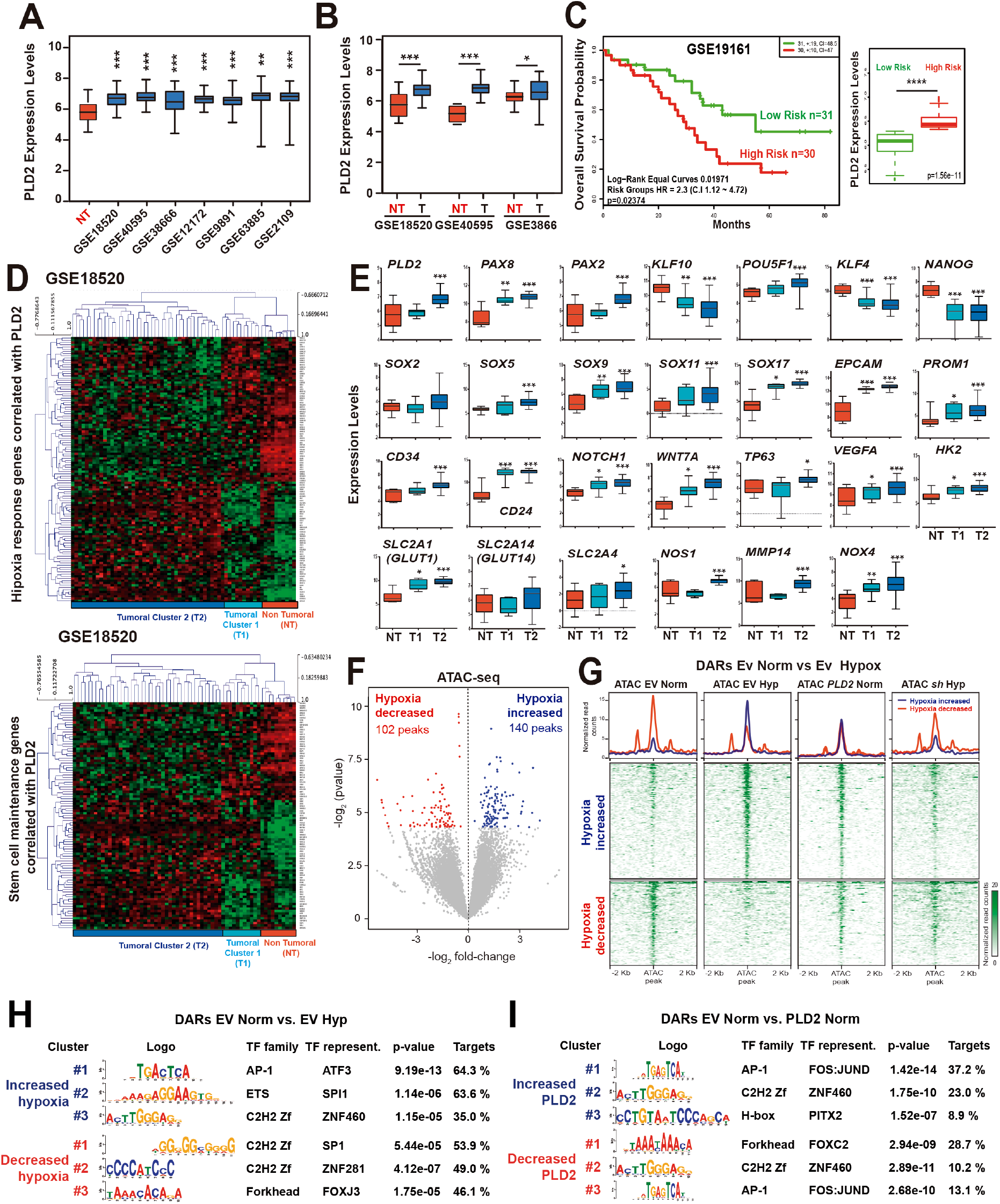
High PLD2 expression in OC patients leads to the transcriptomic rewiring of stemness genes and epigenomic alterations. (**A**) *PLD2* expression in the OC patient databases GSE18520, GSE40595, GSE38666, GSE12172, GSE9891, GSE63885 and GSE2109. (B) *PLD2* expression in the ovarian cancer patient databases GSE18520, GSE40595, and GSE38666. In **(C)** and **(D),** the box plots show the *PLD2* expression levels in ovarian tumor tissue (blue) or nontumor tissue (red) from patients. **(C)** Kaplan–Meier plot showing the overall survival of patients with high (red) or low (green) PLD2 expression levels in the OC patient database GSE19161. The data were analysed with a log-rank test, and the associated P values are shown in the graph. For A to C, expression levels are shown as log_2_ transformed values from the R2 database. **(D)** Heatmaps showing the expression z scores of stem-associated genes or hypoxia response genes whose expression was correlated with *PLD2* in the GSE18520 OC patient database. **(E)** Expression levels of *PLD2* and selected stem-associated or hypoxia response genes correlated with *PLD2* in the OC patient database GSE18520. Box plots showing gene expression in patients in tumoral Cluster 1 (T1; light blue), tumoral Cluster 2 (T2; dark blue), or nontumor tissue (NT; red). Box plots representing the centreline, median; box limits, 25th and 75th percentiles. For A, B, C and E, data were compared using Student’s t tests. Asterisks indicate statistical significance with respect to NT tissue. *p < 0.05; **p < 0.01; ***p < 0.001. **(F)** Differential analyses of accessibility between OC cells carrying Ev under normoxic or hypoxic conditions from ATAC-seq data (n = 2 biological replicates per condition). The log2 normalized read counts of peaks versus the log2-fold-change of accessibility are plotted. Peaks showing a statistically significant change (P value < 0.05) are highlighted in blue (hypoxia increased peaks) or red (normoxia increased peaks). **(G)** Heatmaps plotting normalized ATAC-seq signals at differentially accessible regions (DARs) from (f) in OC cells carrying Ev or expressing *PLD2* under normoxia and carrying Ev or expressing *shPLD2* under hypoxia. **(H-I)** Motif enrichment analyses of the increased and decreased ATAC peaks in OC cells carrying Ev in hypoxia vs. normoxia (h) and +/-*PLD2* expression in normoxia (i). The three motifs with the lowest p values are shown in each case.

Finally, we extended the gene expression analyses to EMT genes using TaqMan Arrays and found that while either hypoxia or *PLD2* overexpression in normoxia led to an increase in the expression of many of these genes, *PLD2* depletion was unable to suppress such an increase, a result further validated by RT‒qPCR of particular EMT genes, and hierarchical clustering of the samples did not exhibit the pattern observed in stemness genes (Supplementary Figure S6A-C). Consistently, invasiveness assays using Boyden’s chamber showed that both *PLD2* overexpression and hypoxia were able to increase invasion, but the *PLD2* depletion did not have any effect (Figure 3F and Supplementary Figure S6D). These results indicate that while the increased expression of stemness genes induced by hypoxia relies on *PLD2* overexpression, this is not the case for EMT genes and suggests that PLD2 is a specific mediator of the increase in CSCs induced by hypoxia in OC cells.

### Hypoxia-mediated reprogramming to pluripotent stem cells is dependent on PLD2

We aimed to obtain additional evidence of the contribution of PLD2 to dedifferentiation or reprogramming events mediated by hypoxia that may generate ovarian CSCs from normal OC cells. Therefore, we performed reprogramming experiments of mouse embryonic fibroblasts (MEFs) to induce pluripotent stem cells (iPSCs) in normoxia and hypoxia and alteration of the *PLD2* expression levels (overexpression and depletion). We used a previously published protocol ^50^ in which MEFs were infected using a HEK293T cell-derived virus that provides OSKM genes and Nanog reporter retroviruses and then cocultured on SNL feeder cells that produce LIF. Then, the samples were incubated with or without hypoxia for 7 days, and the efficiency of iPSC generation was measured for an additional 5 days. Cell reprogramming and the acquisition of pluripotency were assessed by colony morphology, alkaline phosphatase and *nanog* promoter-driven GFP expression analyses (Figure 3G) to assess the effect of PLD2 and hypoxia on the efficiency of the reprogramming process and the acquisition of stem cell-like properties. Using this protocol, we found that, as expected, hypoxia led to a significant increase in the generation of iPSCs (Figure 3H). Furthermore, we found that the *PLD2* overexpression in normoxia provoked a similar increase in iPSC generation, consistent with its effect on the generation of CSCs, and that the combination of both hypoxia and *PLD2* overexpression further increased iPSC formation (Figure 3H), confirming that high *PLD2* expression leads to dedifferentiation processes. Finally, *PLD2* depletion in hypoxia suppressed the increased iPSC production induced by hypoxia, and this was suppressed by recovering *PLD2* expression in *PLD2*-depleted cells (Figure 3H), suggesting that PLD2 is an important mediator in the activation of pluripotency by hypoxic conditions.

### PLD2 is overexpressed in OC patients and leads to a rewiring of the hypoxia and stemness transcriptional programs

Thus far, we showed that hypoxia induces the overexpression of *PLD2* mediated by HIF-1α, which, in turn, leads to increased CSC features in OC. Then, we wondered whether *PLD2* was indeed overexpressed in OC patients. Therefore, we analysed the *PLD2* expression levels in 7 OC patient databases using the R2 platform and found that *PLD2* expression was significantly higher in OC patients than in non-tumoral ovarian tissue (Figure 4A). This result was confirmed by a comparison between patients and control individuals in 3 databases containing their own controls (GSE18520, GSE4595 and GSE3866) (Figure 4B). Indeed, gain or amplification of the *PLD2* gene in approximately 10% of OC patients was also detected in the TCGA database (Supplementary Figure S7A). Next, we wondered whether *PLD2* expression was associated with patient survival. Therefore, we analysed 4 OC databases with available overall survival (OS) data (GSE13876, GSE19161, GSE23554 and GSE31245), and we first separated the patients into low-risk and high-risk groups based on OS. The difference in OS between the groups was statistically significant only in one database, GSE19161 (Figure 4C); however, the expression levels of *PLD2* were significantly higher in the high-risk group than in the low-risk group in the 4 databases (Figure 4C; Supplementary Figure S7B). These data indicate that *PLD2* is commonly overexpressed in OC patients and may be associated with decreased patient survival.

To determine whether *PLD2* expression in OC patients is related to hypoxia and stemness, we analysed the expression of genes related to these functions in the three OC databases with available expression data from control individuals. First, we selected the genes annotated to the Gene Ontology (GO) term “Response to Hypoxia” or “Stem cell maintenance” whose expression was significantly correlated with that of *PLD2* in OC patients (p<0.05; r>0.2 or <-0.2). Then, we performed hierarchical clustering of patient and control individuals based on the expression levels of these genes (Figure 4D; Supplementary Figure S8). In the GSE18520 database, the clustering clearly separated the control individuals (‘non-tumoral’, NT) and a reduced group of patients who we termed ‘Tumoral Cluster 1’ (T1) from most patients who clustered in what we termed ‘Tumoral Cluster 2’ (T2) (Figure 4D). The patients at T1 showed a transcriptional profile of hypoxia-related genes more similar to NT and clearly different from T2. In contrast, the clustering in the GSE4095 and GSE38666 databases clearly separated clusters of NT and tumoral (T) individuals, who showed distinct transcriptional profiles of hypoxia-related genes (Supplementary Figure S8). These results suggest that there is a switch in the expression of hypoxia-related genes in OC tumors compared with healthy ovaries.

Next, we repeated the hierarchical clustering with the genes annotated to the GO term “Stem cell maintenance” whose expression was significantly correlated with that of *PLD2* in OC patients (p<0.05; r>0.2 or <-0.2). Surprisingly, this clustering based on stem-related genes separated the patients and control individuals into the same clusters as the hypoxia-related genes (Figure 4D; Supplementary Figure S8), suggesting a connection between both groups of genes that supports the model of CSC generation induced by hypoxia. Then, we plotted the expression levels of *PLD2* in the 3 clusters obtained from the GSE18520 database and found that *PLD2* exhibited significantly increased expression in Cluster T2 compared with that in Cluster NT, while Cluster T1 showed similar levels to NT (Figure 4E). This finding suggests that a connection exists among PLD2, the hypoxia response and stemness since *PLD2* expression is misregulated only in patients showing transcriptional profiles related to hypoxia and stemness highly different from healthy controls. In addition, we checked the expression of stem-associated and hypoxia-related genes in these clusters. As shown in Figure 4E, the expression of *PAX8*, a well-known OC marker, was significantly increased in both patient Clusters T1 and T2, similar to other stemness genes, such as *SOX9*, *SOX17*, *EPCAM*, *PROM1*, *CD24*, *NOTCH1* and *WNT7A*. Other genes in this group showed a significant increase in expression only in Cluster T2, coinciding with higher *PLD2* levels, including *PAX2*, *POU5F1* (*OCT4*), *SOX5*, *SOX11*, *CD34* and *TP63*, while the others were unaffected or even significantly reduced, such as *KLF4*, *NANOG* or *SOX2*, although the latter showed a nonsignificant increase in Cluster T2 (Figure 4E). This finding suggests that there is an OC stemness signature in patients who may be stronger with higher *PLD2* expression. However, some hypoxia-related genes showed a significant increase in expression in both Clusters T1 and T2, including *VEGFA*, *SLC2A1* (*GLUT1*), *HK2* and *NOX4*, or only in T2, including *SLC2A4*, *NOS1* and *MMP14*, and a nonsignificant increase in *SLC2A14* (*GLUT14*) (Figure 4E). Similar results were obtained in the NT and T clusters in the GSE4095 and GSE38666 databases (Supplementary Figure S8). Altogether, these results suggest that there is transcriptional rewiring of the expression of hypoxia– and stem-related genes in OC patients with *PLD2* overexpression.

### Hypoxia and PLD2 modify the epigenomic landscape of OC cells

To analyse whether the effect of hypoxia and PLD2 on the expression of stemness-related genes was caused by changes in the activity of the CREs controlling these genes, we performed ATAC-seq experiments in SKOV3 cells under normoxia and hypoxia and altered *PLD2* expression under normoxia (*PLD2* overexpression) and hypoxia (*PLD2* depletion). We computationally called peaks in normoxia and hypoxia and compared both conditions by a differential accessibility analysis; we detected 140 and 102 peaks with increased or decreased accessibility in hypoxia, respectively (Figure 4F). The heatmaps and aggregate profiles of these differentially accessible regions (DARs) showed that the peaks with increased accessibility in hypoxia were also more open upon *PLD2* overexpression in normoxia, although to a lower extent, and vice versa, with the peaks showing decreased accessibility in hypoxia (Figure 4G), suggesting that *PLD2* overexpression in normoxia has a similar effect on chromatin accessibility as hypoxia. Indeed, the DARs of EV-versus *PLD2*-overexpressing cells under normoxia showed similar changes in accessibility under hypoxia, reinforcing the previous idea (Supplementary Figure S9A-B). Both DARs in normoxia versus hypoxia and control versus *PLD2* overexpression were associated with genes enriched in Gene Ontology terms related to the response to hypoxia or well-known functions of *PLD2*, respectively (Supplementary Figure S9C-D). Finally, DARs with increased or decreased accessibility in hypoxia control cells showed decreased or increased accessibility upon *PLD2* depletion, respectively (Figure 4G), and the same occurred with DARs in *PLD2*-overexpressing cells in normoxia (Supplementary Figure S9B), suggesting that PLD2 and hypoxia lead to a similar rewiring of the epigenome in OC cells and that the effect of hypoxia partially depends on *PLD2* expression.

Next, we sought to investigate the possible mechanisms driving the alterations in the chromatin accessibility landscape by hypoxia and *PLD2* overexpression. Therefore, we first performed motif enrichment analyses of DARs. We found that DARs in hypoxia were enriched in the motifs of the AP-1, ETS and C2H2 zinc finger transcription factor families in the increased accessibility sites and the C2H2 zinc finger and fork head families in the decreased accessibility sites (Figure 4H). Similar enrichments were found in DARs upon *PLD2* overexpression (Figure 4I), reinforcing the idea of a similar effect under both conditions. To further elucidate the possible TFs involved in hypoxia and PLD2-mediated epigenomic changes, we estimated differential TF binding among the conditions based on footprints in ATAC-seq data. Using this approach, we confirmed the increased chromatin binding of AP-1 family transcription factors, such as FOS and JUN, consistent with their implication in the response to hypoxia ^51^ (Figure 5A-B). We also found other TF families showing increased TF binding in hypoxia, such as homeobox, paired box or fork head TFs, and the C2H2 zinc finger TF ZBTB32, while TF families with decreased binding in hypoxia included members of the bHLH, NF-Y and ETS families, among others (Figure 5A-B). When we compared these TFs with those showing increased binding upon *PLD2* overexpression in normoxia, we found that most of these TFs overlapped (Figure 5C). In particular, 25 TF motifs of the AP-1 family showed increased binding under both conditions. Interestingly, these AP-1 and 20 more TF motifs with increased binding in hypoxia showed decreased binding upon *PLD2* depletion (Figure 5C-E), indicating that they are dependent on *PLD2* expression. Altogether, these results suggest a function of the AP-1 family of TFs mediating the alterations in the epigenomic landscape mediated by hypoxia and PLD2.

**Figure 5.**
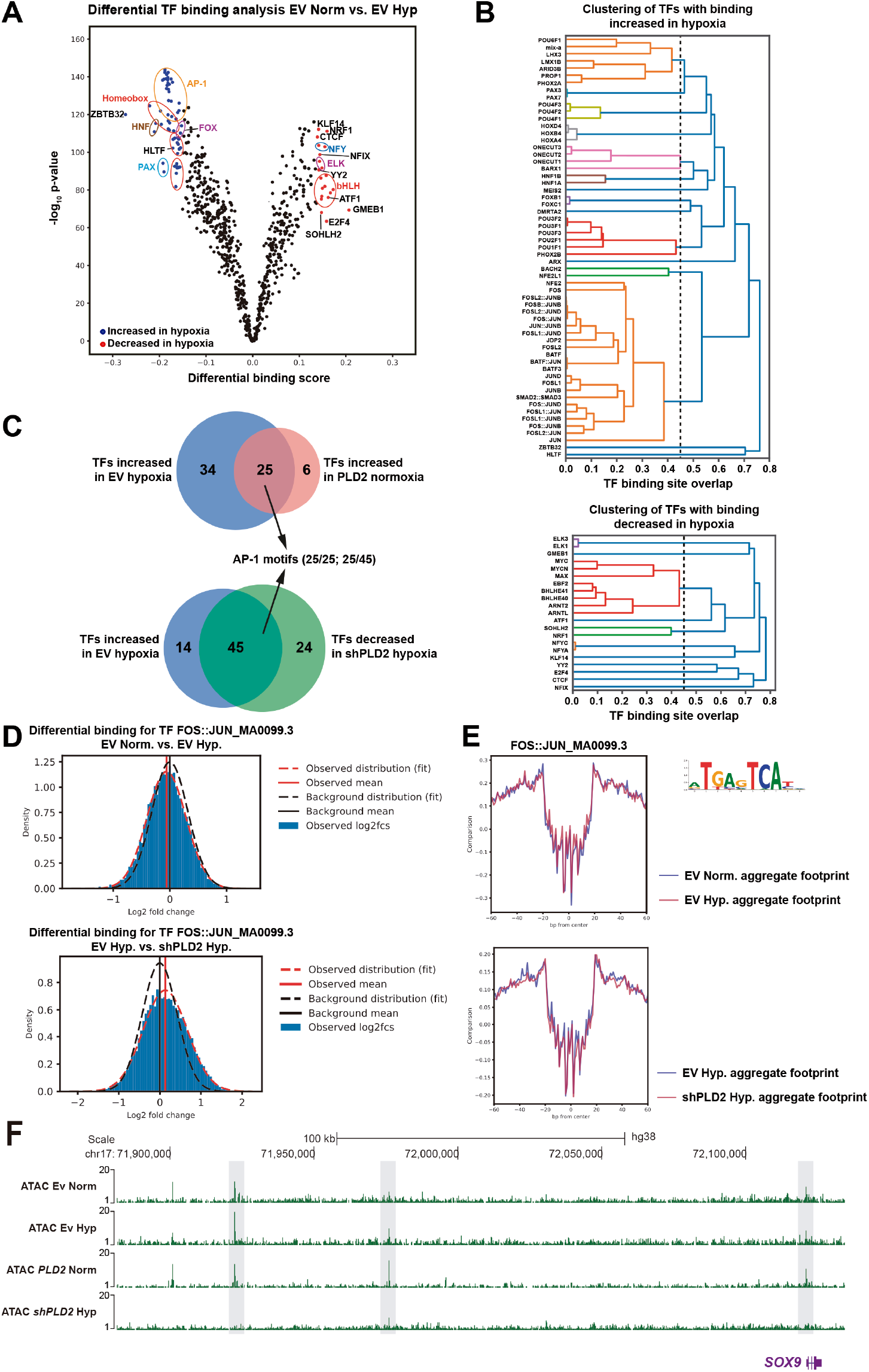
Hypoxia and PLD2 modify the epigenomic landscape of OC cells. (**A**) Differential transcription factor (TF) binding analysis in OC cells carrying Ev under normoxia vs. hypoxia conditions using TOBIAS. Volcano plot showing the differential binding score versus the –log_10_ p value. TF motifs with increased (blue) or decreased (red) binding in hypoxia are highlighted. **(B)** Clustering of TFs with increased (top) or decreased (bottom) binding in hypoxia. **(C)** Venn diagrams plotting the overlap between TFs with increased binding in hypoxia and expressing *PLD2* in normoxia (top) or the overlap between TFs with increased binding in hypoxia and expressing *shPLD2* in hypoxia (bottom). **(D)** Distribution of fold changes in the ATAC peaks containing the motifs FOS::JUN in Ev normoxia vs. Ev hypoxia (top) or in Ev hypoxia vs. *shPLD2* hypoxia**. (E)** Aggregate footprint signal of the peaks containing the motifs in d. **(F)** Tracks with ATAC-seq signals in OC cells carrying Ev in hypoxia or normoxia, expressing *PLD2* in normoxia or expressing *shPLD2* in hypoxia at the *SOX9* locus.

Finally, we analysed whether these alterations in the chromatin accessibility landscape of OC cells could be connected to the elevated expression levels of stemness genes. Therefore, we first selected ATAC peaks falling within the putative regulatory landscapes of the genes associated with stem cell maintenance and proliferation. The clustering of these 3,572 peaks revealed 4 groups with different accessibility levels and behaviours (Supplementary Figure S10A). Among them, Cluster 3 corresponded with peaks with increased accessibility in both hypoxic and *PLD2*-overexpressing cells but decreased accessibility in *PLD2*-depleted cells. Among the stemness genes associated with this cluster, we found genes with increased expression levels in OC cells subjected to hypoxia or *PLD2* overexpression and/or overexpressed in OC patients, such as *SOX9* (Figure 5F), *PROM1*, *JAG1* (functionally connected to *NOTCH1*) or *WNT7A* (Supplementary Figure S10B). These results indicate that stemness genes overexpressed under hypoxic and *PLD2*-overexpressing conditions are connected to alterations in the accessibility of their associated CREs, a part of which is dependent on *PLD2* expression.

### Overexpression of *PLD2* leads to chemotherapy resistance in ovarian tumors

Since we showed that *PLD2* overexpression leads to an increase in CSC-like cells in OC and CSCs were previously proposed to be responsible for chemotherapy resistance and tumor relapse, we wondered whether *PLD2* overexpression could cause resistance to conventional therapy in ovarian tumors. Therefore, we first analysed the expression levels of *PLD2* in our own cohort of OC patients. The immunohistochemistry analyses showed that the *PLD2* protein levels were higher in tumors than in healthy tissue (Figure 6A), and RT‒qPCR revealed that *PLD2* mRNA was significantly more abundant in OC patients than in control non-tumoral samples (Figure 6B), thus confirming the results observed in the transcriptomic databases (Figure 4A-B). Then, we separated our patient samples into those who were sensitive or resistant to platinum-based chemotherapy (without or with tumor relapse within the next 6 months after chemotherapy, respectively) and analysed the *PLD2* expression levels. Importantly, we found that the resistant patients showed significantly higher expression of *PLD2* than the sensitive patients (Figure 6C), suggesting that *PLD2* overexpression may contribute to resistance to platinum-based therapy. The resistant patients in our cohort showed reduced OS and PFS (Figure 6D-E), consistent with the reduced survival of patients with high *PLD2* expression (Figure 6C).

**Figure 6.**
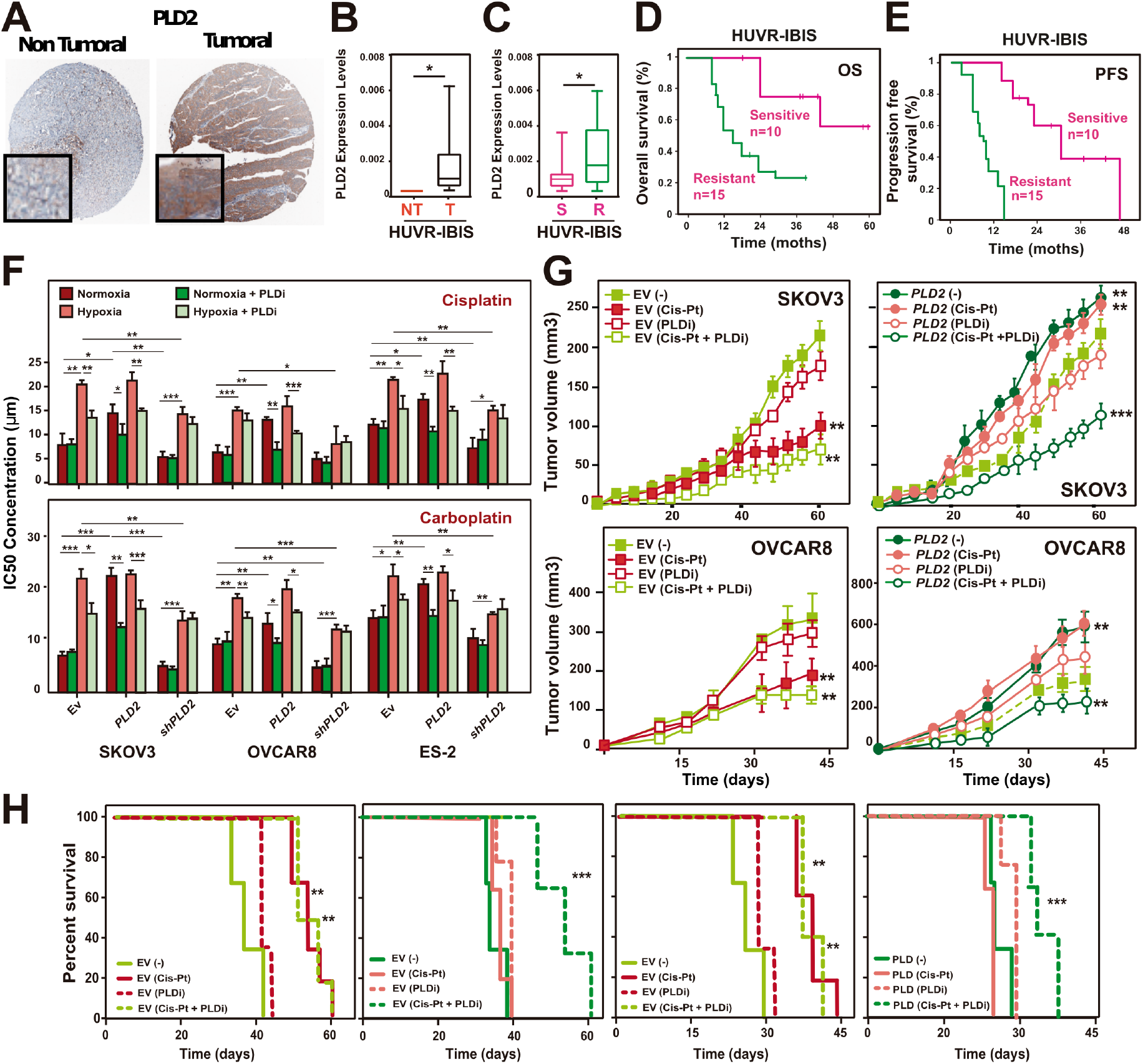
*PLD2* overexpression is associated with resistance to treatment in ovarian cancer. (**A**) Representative images of PLD2 immunostaining in ovarian cancer and non-tumoral samples. Scale bars, 50 µm. **(B)** *PLD2* expression in ovarian patients from the HUVR-IBIS database. Box plots showing the *PLD2* expression levels in ovarian tumor tissue (black) or nontumor tissue (red) from patients. Box plots representing the centreline, median; box limits, 25th and 75th percentiles; whiskers, minimum and maximum values. **(C)** Analysis of *PLD2* expression by RT‒qPCR in a cohort of OC patients who were sensitive (S; pink) or resistant (R; green) to platinum treatment (HUVR-IBIS). For B and C, the mRNA expression was calculated as 2^-ΔCt^ relative to the *ACTB* gene. **(D-E)** Kaplan‒Meier plots showing overall or progression-free survival in patients who were sensitive (pink) or resistant (green) to platinum treatment in the HUVR-IBIS cohort. **(F)** Determination of the IC_50_ of cis-platinum and carboplatin in ES-2, SKOV3 and OVCAR8 cells overexpressing *PLD2* or *shPLD2* or carrying EV in normoxia and hypoxia, in combination or without the PLD inhibitor (PLDi) FIPI in ES-2, SKOV3 and OVCAR8 cells overexpressing *shPLD2* or *PLD2* or carrying EV in normoxia or hypoxia. **(G-H)** Determination of the tumor volume (G) and survival (H) after treatment with saline, cisplatin, PLDi or both in xenografts of SKOV3 cells expressing *shPLD2, PLD2* or EV. A minimum of three biological were performed per each experiment. The data were compared using Student’s t tests. For F to H, asterisks indicate statistical significance with respect to Ev carrying cells or xenografts, unless indicated by horizontal lines. *p < 0.05; **p < 0.01; ***p < 0.001.

Next, we analysed the effect of *PLD2* expression on resistance to platinum compounds in OC cells *in vitro*. We first found that *PLD2* expression in public data from gynaecological cancer cells exposed to cisplatin treatment showed little changes among control and treated cells, and only some of the analysed CSC genes showed notable changes in particular cell lines (Supplementary Figure S11). Thus, cells overexpressing or depleted of *PLD2* were treated with increasing concentrations of cisplatin and carboplatin, and the IC_50_ values were calculated in each case. We found that *PLD2* overexpression led to a significant increase in the IC_50_ values, while the *PLD2* depletion led to only a weak nonsignificant reduction (Figure 6F). This finding suggests that higher *PLD2* expression causes resistance to platinum-based compounds. We repeated these experiments under hypoxic conditions and found that hypoxia also led to increased IC_50_ values (Figure 6F). However, the *PLD2* depletion reduced the increase in the IC_50_ values induced by hypoxia, suggesting that increased resistance to cisplatin and carboplatin in OC cells under hypoxic conditions relies on *PLD2* expression.

Finally, we performed *in vivo* analyses to validate our findings by establishing xenograft models from SKOV3 and OVCAR8 cells overexpressing or depleted of *PLD2* and analysing tumor growth upon treatment with cisplatin. The control tumors from cells transfected with the empty vector were sensitive to the cisplatin treatment, significantly reducing tumor growth in xenografts from both SKOV3 and OVCAR8 cells (Figure 6G). However, the tumors overexpressing *PLD2* showed increased tumor growth that was not reduced upon the cisplatin treatment, indicating both a higher aggressiveness of *PLD2*-overexpressing tumors and the resistance of these tumors to cisplatin. However, the *PLD2* depletion resulted in significantly reduced tumor growth that was further reduced upon the cisplatin treatment (Figure 6G). Importantly, the cisplatin treatment in the control tumors led to increased survival, while the *PLD2*-overexpressing tumors did not exhibit improved survival, and the *PLD2*-depleted tumors exhibited increased survival independent of the cisplatin treatment (Figure 6H). These results indicate that the overexpression of *PLD2* causes resistance to platinum-based chemotherapy in OC tumors.

### Combination treatment with cisplatin and PLD inhibitor suppresses chemotherapy resistance

Finally, we wondered whether the increased therapy resistance to platinum-based compounds induced by *PLD2* overexpression and hypoxia could be suppressed by the pharmacological inhibition of PLD2. For this, we used the PLD inhibitor (PLDi) 5-Fluoro-2-indolyl des-chlorohalopemide (FIPI), which inhibits PLD2 catalytic activity ^52^. First, we tested this possibility *in vitro* by calculating the IC_50_ in OC cells treated with cisplatin, PLDi and their combination in normoxia and hypoxia with altered levels of *PLD2*. Interestingly, we found that the higher IC_50_ of cisplatin in the OC cells overexpressing *PLD2* or in hypoxia was suppressed by the use of PLDi (Figure 6F). Next, we validated these results *in vivo* by establishing xenografts of OC cells expressing EV or *PLD2* and treating the mice with cisplatin, PLDi or their combination. We found that the increased tumor growth provoked by *PLD2* overexpression was reduced upon the treatment with PLDi (Figure 6G). Moreover, although the treatment with cisplatin did not reduce the higher tumor growth induced by *PLD2* overexpression, its combination with PLDi led to a significant reduction in tumor growth that was stronger than that following the treatment with PLDi alone (Figure 6G). This finding was confirmed in xenografts from two OC cell lines (OVCAR8 and SKOV3) and led to an increase in survival (Figure 6H). Altogether, these results indicate that the chemotherapy resistance to cisplatin caused by *PLD2* overexpression can be overcome by the pharmacological inhibition of PLD2, suggesting that combined treatment with cisplatin and PLDi is a promising alternative treatment for patients with high *PLD2* expression levels.

## DISCUSSION

Using *in vitro* and *in vivo* models of OC, patient samples and public databases, we show that the HIF-1α-PLD2 axis is a major player in the chemoresistance of OC by promoting the generation of CSCs under hypoxic conditions. On the one hand, *PLD2* expression is regulated by HIF-1α through HREs in its promoter and a hypoxia-specific enhancer; on the other hand, *PLD2* expression is required for the full transcriptional and epigenomic rewiring promoted by hypoxia as evidenced using gene expression and chromatin accessibility analyses. In particular, the expression of stem-related genes, the opening of enhancers in the vicinity of these genes, the generation of CSCs induced by hypoxia, and the reprogramming of normal cells to iPSCs rely on normal *PLD2* expression levels. These findings indicate that PLD2 is a major player in the response to hypoxia in cancer cells that leads to increased stem cell properties resulting in higher chemoresistance.

Hypoxia is a feature of the tumor microenvironment in regions with low oxygen supply that is known to increase the stemness features of cancer cells in several types of cancer ^18–22, 53^, including OC in which hypoxia has been shown to increase the stem-like properties of cancer cells ^27^. HIF-ARNT can regulate the expression of many genes that promote the hypoxic response ^2, 3^. Therefore, HIF factors may exert their effect of increasing CSCs via multiple mechanisms. In OC, HIF factors contribute to the upregulation of pluripotency factor genes, such as *SOX2* or *OCT3/4* ^27, 54^, proliferation pathways, such as Notch or Wnt ^54, 55^, or epigenetic modulation by affecting chromatin modifiers, such as SIRT1 ^28^. Here, we show that the OC cell lines ES-2, SKOV3 and OVCAR8 under hypoxic conditions show upregulated expression of *PLD2*, which encodes phospholipase D2, in these cells in a Hif-1α-dependent manner (Figure 1), as recently reported in colon cancer ^42^. Using DNA binding motif analyses and reporter assays, we also demonstrate that *PLD2* expression is regulated at the transcriptional level by HREs located within the *PLD2* promoter and a hypoxia-specific enhancer that activates *PLD2* transcription in hypoxia, thus providing a mechanistic explanation of Hif-1α-mediated *PLD2* overexpression. Therefore, it is likely that the HIf-1α-PLD2 axis works in other solid tumors under hypoxic conditions.

Importantly, *PLD2* depletion partially suppresses the effect induced by hypoxia (Figure 3), suggesting that PLD2 is an important mediator of hypoxia-induced stemness. This finding was also corroborated by the reprogramming experiments of MEFs to iPSCs in which PLD2 was required for the increased reprogramming induced by hypoxia. Indeed, PLD2 might also potentiate the response to hypoxia by a positive feedback loop. This hypothesis can be supported since the combination of hypoxia and *PLD2* overexpression additively enhanced the stem properties of OC cells (Figure 3). The activation of HIF-1α expression or activity by PLD2 has been reported in endothelial, glioma and renal cancer cells ^43–45^, but there is evidence of the opposite effect in HEK293 cells ^46^, suggesting that this feedback loop may work in specific contexts or cell types. Nevertheless, whether the effect of PLD2 on stemness is an autocrine or a paracrine effect remains to be elucidated. In this regard, we previously showed that exosomes from *PLD2*-overexpressing colorectal cancer cells induced senescence in stromal fibroblasts ^41^, which, in turn, induced WNT pathway activation and increased stemness in tumor cells. In OC, hypoxia-induced exosomes have been involved in increased tumorigenic properties and chemoresistance by several mechanisms. These include exosome-containing oncogenic proteins, such as STAT3 and FAS ^56^, microRNAs that alter tumor-associated macrophages ^57, 58^, and plasma gelsolin, which induces the conversion of chemosensitive OC cells to chemoresistant cells ^59^. Finally, a role of PLD1 and PLD2 inducing exosome secretion in OC cells has also been recently proposed ^60^, suggesting that PLD2 may also influence the tumor microenvironment in OC.

The expression of *PLD2* is elevated in several cancer types ^35–41^, and here, using public patient databases and our cohort of patients, we show that this is also the case in OC in which it may be related to decreased OS (Figure 4). Furthermore, consistent with the results in the OC cell lines, the clustering analyses of OC patient gene expression revealed that *PLD2* expression is correlated with the rewiring of transcriptomic programs of the response to hypoxia and stem cell maintenance. Indeed, we found two different clusters of patients, one of which showed gene expression patterns more different from healthy controls coinciding with higher *PLD2* expression. Although the expression of the OC marker *PAX8* and other stemness– and hypoxia-related genes was enhanced in both clusters, other markers showed increased expression only in the patient cluster with high *PLD2* levels, suggesting that PLD2 may promote stemness through specific genes or pathways as we show in *SOX2*, *SOX9* and *NOTCH1,* but not *SOX17*. These data indicate that a correlation exists between *PLD2* expression and highly altered transcriptomic programs of the response to hypoxia and stemness. In addition, we provide evidence that PLD2 is important for the alteration in the epigenomic landscape provoked by hypoxia since both hypoxia and *PLD2* overexpression in normoxia lead to similar alterations in chromatin accessibility that are counteracted by *PLD2* depletion, including the opening of enhancers in the proximity of genes related to the stem fate, which we show were upregulated (Figure 4). These changes likely occur through the activation of CREs bound by the AP-1 family of TFs (Figure 5). In a previous study analysing OC tumors, solid metastasis and effusions, higher *PLD2* expression was described in effusions rather than solid tumors and metastasis ^47^. This finding is consistent with our findings since OC effusions, most of which are peritoneal, are characterized by low oxygen levels and a high content in CSCs ^26^. Therefore, the hypoxia-PLD2 axis seems to play a major role in the generation of ovarian CSCs.

In OC, several mechanisms have been described to promote chemotherapy resistance under hypoxic conditions, including the upregulation of the *ABCG2* transporter gene, which increases drug efflux ^29, 30^, *c-KIT* overexpression ^55^ and high cysteine levels ^61^. CSCs are responsible for chemotherapy resistance ^14, 15^, and ovarian CSCs were identified sixteen years ago and reported to be chemoresistant ^62, 63^. In agreement with this idea, we previously found several markers linking ovarian CSCs and chemoresistance ^64, 65^. Here, using OC cells and xenograft models, we show that the overexpression of *PLD2* leads to resistance to platinum-derived compounds, including cisplatin and carboplatin (Figure 6). Moreover, *PLD2* expression is higher in patients resistant to platinum-based chemotherapy than in sensitive patients, confirming our results in the cell lines and mouse models. How PLD2 provokes such resistance is an intriguing issue, although it is likely that its enzymatic product PA, an important molecule acting as a second messenger in multiple cellular functions ^66^, might play some relevant role in avoiding chemotherapy-induced cell damage. Indeed, we previously showed that PA administration has similar effects as *PLD2* overexpression ^41^. This is particularly relevant in ovarian tumors, which are typically treated with platinum-based compounds and show high rates of chemoresistance, with frequent metastasis in hypoxic environments, such as abdominal ascites. Therefore, we propose an alternative treatment based on a combination of cisplatin and the pharmacological inhibition of PLD2. Our *in vitro* and *in vivo* results demonstrate that this combined treatment may be useful for patients with high expression levels of *PLD2* who are resistant to conventional therapy with cisplatin alone.

In summary, our findings suggest a model in which hypoxia leads to the transcriptional overexpression of *PLD2* in OC, which, in turn, generates PA and induces the generation of chemoresistant ovarian CSCs. Thus, our work highlights the importance of the HIF-1α-PLD2 axis for CSC generation and chemoresistance in OC cells. However, hypoxia induces many stemness signals depending on the tissue, mostly mediated by HIF-1α, but the transcriptional changes induced by hypoxia-inducible factors (HIFs) also activate other stemness signals that might activate pluripotency and stemness in tumor cells ^18, 20, 22, 53^. Since part of the stemness induced by hypoxia in OC is dependent on PLD2 activation, how is stemness signalling rewired in these tumors? It is possible that it has a specific effect on ovarian tissue. It is also possible that PLD2 may account for the activation of some secondary stemness signalling. Most likely, there is some partial redundancy among the different stemness signalling pathways, increasing the robustness of the wiring in tumor cells. Nevertheless, more work is needed to expand our knowledge of CSCs in the tumorigenic process.

## METHODS

### Cell culture

Cells were cultured according to the manufacturer’s recommended procedures in McCoy (ES-2 line) or RPMI (SKOV3 and OVCAR8 lines) and incubated at 37 °C in 5% CO_2_ in a humidified atmosphere.

### Gene transfer

The gene transfer was performed as previously described ^65^. The PLD2 overexpression plasmid was described in ^41^. The shRNA against *PLD2* was provided by Origene.

### Proliferation assay

The proliferation assay was performed as previously described ^65^.

### Cytotoxic assay

ES-2, SKOV3 or OVCAR8 cells were seeded and then treated with platinum drugs and/or the PLD inhibitor 5-Fluoro-2-indolyl des-chlorohalopemide (FIPI) at 300nM concentration 24 hours later. After 96 hours, cells were stained with 0.5% crystal violet. Then, the crystal violet was solubilized in 20% acetic acid and quantified at 595 nm absorbance to measure the cell viability.

### Maintenance of mouse colonies

All experiments involving animals received expressed approval from the IBIS/HUVR Ethical Committee for the Care and Health of Animals. The mice were maintained in the IBIS animal facility according to the facility guidelines, which are based on the Real Decreto 53/2013 and were sacrificed by CO_2_ inhalation either using a planned procedure or as a human endpoint when the animals showed significant signs of illness.

### Colony formation assay and clonal heterogeneity analysis

This analysis was performed as previously described in ^65^. Briefly, in total, 10^3^ cells were seeded onto 10 cm plates, and every condition was evaluated in triplicate. The medium was replaced every 3 days for 12 days, and the colonies were fixed, stained and counted. The values are expressed as the number of observed colonies among the 10^3^ seeded cells. To analyze the clonal heterogeneity, 10^2^ random colonies were classified in triplicate as having the following phenotypes: holoclone, meroclone and paraclone ^67^, which are considered stem cells, transit-amplifying cells and differentiated cells, respectively ^49^.

### Sphere-forming assay

In total, 2×10^3^ cells were resuspended in 1 ml of complete MammoCultTM Basal Medium (Stemcell Tech) and seeded in ultralow attachment plates. The cultures were imaged, the tumorspheres were counted, and their diameters were quantified using CellSenseDimension software on Days 2, 3 and 4.

### Western blot analyses and immunofluorescence

were performed according to standard procedures. Information of antibodies and dilutions is shown in Supplementary Table S1.

### RT–qPCR

The total RNA was isolated using an RNeasy kit (Qiagen), and cDNA was generated from 1 µg of RNA with MultiScribe Reverse Transcriptase (Applied Biosystems). qPCR was performed using a TaqMan Assay (Applied Biosystems) with probes. The relative mRNA expression was calculated as 2^-ΔCt^ relative to the *ACTB* gene. Information of probes is shown in Supplementary Table S1.

### Fluorescence-activated cell sorting

For FACS staining, live cells were incubated with antibodies for 30 minutes at dilutions specified in the manufacturer’s protocols.

### iPSCs protocol

Briefly, mouse embryonic fibroblasts (MEFs) were infected with 4 different retroviral vectors encoding the Yamanaka factors (pMXs-Oct3/4, pMXs-Sox2, pMXs-Klf4, and pMXs-cMyc), an additional lentiviral vector that expresses GFP in cells where the Nanog promoter/enhancer is active (mNanog-pGreenZeo), and the corresponding plasmid overexpressing PLD2, carrying shPLD2 or carrying Ev. Seventy-two hours after the infection, MEFs were seeded on top of an SNL feeder layer in the presence of ES media, and the media was renewed every 24 h. Cell reprogramming and the acquisition of pluripotency were assessed by colony morphology, GFP expression and alkaline phosphatase activity assays to assess the effect of the gene of interest/condition of interest* on the efficiency of the reprogramming process and the acquisition of stem cell-like properties.

### ATAC-seq

ATAC-seq assays were performed using standard protocols ^68, 69^, with minor modifications. Briefly, 70,000 ovarian cancer cells overexpressing PLD2 carrying shPLD2 or carrying Ev growing under normoxic or hypoxic conditions were collected by centrifugation for 5 min at 500 g 4 °C. The supernatant was removed, and the cells were washed with PBS. Then, the cells were lysed in 50 µl of lysis buffer (10 mM Tris-HCl pH 7.4, 10 mM NaCl, 3 mM MgCl_2_, 0.1% NP-40, 1× Roche Complete protease inhibitors cocktail) by pipetting up and down. The whole cell lysate was used for TAGmentation, which was centrifuged for 10 min at 500 g 4 °C, resuspended in 50 µl of the Transposition Reaction containing 2.5 µl of Tn5 enzyme and TAGmentation Buffer (10 mM Tris-HCl pH 8.0, 5 mM MgCl2, 10% w/v dimethylformamide), and incubated for 30 min at 37 °C. Immediately after TAGmentation, DNA was purified using a Minelute PCR Purification Kit (Qiagen) and eluted in 20 µl. Libraries were generated by PCR amplification using NEBNext High-Fidelity 2X PCR Master Mix (NEB). The resulting libraries were multiplexed and sequenced in a HiSeq 4000 paired-end lane, producing 100 M 49-bp paired-end reads per sample.

### Quantification and statistical analysis

All statistical analyses were performed using GraphPad Prism 4. The distribution of the quantitative variables among different study groups was assessed using parametric (Student’s *t test*) or nonparametric (Kruskal–Wallis or Mann– Whitney) tests as appropriate. The experiments were performed a minimum of three times and were performed in independent triplicates each time. The survival data from the patient databases were analyzed by a log-rank Mantel‒Cox statistical test.

### Analyses of cancer patient databases

We performed meta-analyses using the R2 Genomics analysis and visualization platform (http://hgserver1.amc.nl) to analyze the *PLD2* expression levels in tumor and nontumor ovarian samples from the databases. The statistical significance of the tumor versus normal samples was assessed (P<0.05). Patient survival was analyzed using an R2 Genomics analysis and visualization platform (http://hgserver1.amc.nl), which was developed by the Department of Oncogenomics of the Academic Medical Center (Amsterdam, Netherlands). Kaplan‒Meier plots showing patient survival were generated using the databases with available survival data with the scan method, which searches for the optimum survival cut-off based on statistical analyses (log-rank test), thereby identifying the most significant expression cut-off.

### ATAC-seq data analyses

ATAC-seq reads were aligned to the GRCh38 (hg38) human genome assembly using Bowtie2 2.3.5 ^70^, and pairs separated by more than 2 kb were removed. For ATAC-seq, the Tn5 cutting site was determined as position –4 (minus strand) or +5 (plus strand) from each read start, and this position was extended 5 bp in both directions. Reads below 150 bp were considered nucleosome free. The conversion of the SAM alignment files to BAM was performed using SAMtools 1.9 ^71^. The conversion of BAM to BED files and peak analyses, such as overlaps or merges, were carried out using the Bedtools 2.29.2 suite ^72^. The conversion of BED to BigWig files was performed using the genomecov tool from Bedtools and the wigToBigWig utility from UCSC ^73^. For ATAC-seq, peaks were called using the MACS2 2.1.1.20160309 algorithm ^74^ with an FDR < 0.05 for each replicate and merged into a single pool of peaks that was used to calculate the differentially accessible sites with the DESeq2 1.18.1 package in R 3.4.3 ^75^; a P value < 0.05 was set as the cut-off for statistical significance of the differential accessibility. For visualization purposes, reads were extended 100 bp for ATAC-seq. For the data comparison, all ATAC-seq experiments used were normalized using reads falling into peaks to counteract differences in background levels between experiments and replicates ^76^.

Heatmaps, average profiles and k-means clustering of the ATAC-seq data were generated using the computeMatrix and plotHeatmap tools from the Deeptools 3.5 toolkit ^77^. The TF motif enrichment was calculated using XSTREME ^78^ with the standard parameters. For the gene assignment to ATAC peaks, we used the GREAT 3.0.0 tool ^79^, with the basal plus extension association rule and the standard parameters (5 kb upstream, 1 kb downstream, and 1 Mb maximum extension). For the footprinting analyses, we used TOBIAS 0.12.9 ^80^. First, we performed bias correction using ATACorrect and calculated the footprint scores with ScoreBigwig, both with the standard parameters. Then, we used BINDetect to determine the differential TF binding of all vertebrate motifs in the JASPAR database ^81^. We considered motifs with a linear fold-change ≥ 15% between conditions differentially bound. Aggregated ATAC-seq signals in the footprints were visualized using PlotAggregate. ATAC-seq data generated in this study is available through the Gene Expression Omnibus (GEO) accession number GSE210599.

### *In vivo* xenograft studies

Tumor growth was assayed following the subcutaneous injection of 4×10^6^ SKOV3 or OVCAR8 cells that were transfected with a plasmid carrying PLD2 or shRNA against *PLD2* in cohorts of five nude mice each that were analyzed weekly. The tumors were measured using callipers. All mice were sacrificed once the growth experiment was completed.

### *In vivo* xenograft treatment

Tumors were harvested when they reached 1500 mm^3^, cut into 2×2×2 mm pieces and reimplanted. Mice were randomly allocated to the drug-treated and control-treated (solvent only) groups, and once the tumor reached 20 mm^3^, the mice received the appropriate treatment for 4 weeks (2 doses/week). The mice were monitored daily for signs of distress and weighed twice a week. The tumor size was measured, and the size was estimated according to the following equation: tumor volume = [length x width^2^]/2. The experiments were terminated when the tumor reached 350 mm^3^ or when the clinical endpoint was reached. The drugs cisplatin and carboplatin were obtained from Pharmacy HUVR and were freshly prepared and administered by intraperitoneal injection. We used higher doses in mice, assuming a 70 kg average weight for humans (125 mg/dose in humans) ^65^. We administered two doses per week as follows: 3.5 mg/kg of cisplatin (equivalent to 7 mg/kg, averaging 25 g body weights of each mouse) with or without 3 mg/kg of FIPI. We did not observe signs of toxicity.

### *In vivo* xenografts from tumorspheres

This assay involved the subcutaneous injection of 1×10^3^ cells grown as tumorspheres into the hind legs of 4-week-old female athymic nude mice. The animals were treated as previously described, examined twice a week, incubated for 4 more weeks, and killed, and the tumors were extracted. The tumors were measured using callipers.

### Patient cohort

The entire procedure was approved by the local ethical committee of the HUVR (CEEA O309-N-15). A cohort of paraffin-embedded tissue samples from 25 patients with ovarian cancer was obtained from the biobank of Hospital Universitario Virgen del Rocío-Instituto de Biomedicina de Sevilla (Sevilla, Spain) for the RNA expression studies and the evaluation of the correlation of the clinicopathological features (see Supplementary Table S2). The samples were obtained from biopsies of patients who were subjected to platinum treatment and who were evaluated for their response according to the RECIST criteria; normal tissue, platinum-resistant tumor samples and platinum-sensitive tumor samples were obtained. The tumor samples were sent to the pathology laboratory for diagnosis and were prepared for storage with formalin fixation and paraffin embedding. The samples were stained with haematoxylin/eosin, and RNA was extracted from the tumor tissue.

## DECLARATIONS

### Ethics approval and consent to participate

All methods were performed in accordance with the relevant guidelines and regulations of the Institute for Biomedical Research of Seville (IBIS) and University Hospital Virgen del Rocio (HUVR). All animal experiments and the entire procedures of the patient cohort were performed according to the experimental protocol approved by HUVR Animals Ethics (CEI 0309-N-15).

### Consent for publication

Written consent for publication was obtained from all patients involved in our study.

### Data availability

The datasets used and/or analysed during the current study are available from the corresponding author upon reasonable request. ATAC-seq data generated in this study is available through the Gene Expression Omnibus (GEO) accession number GSE210599.

### Conflict of interest

The authors declare that they have no other competing financial interests.

### Funding

This research was funded by Grants RTI2018-097455-B-I00 and PID2021-122629OB-I00 funded by MCIN/AEI/10.13039/501100011033 and by “ERDF A way of making Europe”, by the “European Union”. Additional grants from CIBER de Cáncer (CB16/12/00275), from Consejeria de Salud (PI-0397-2017) and Project P18-RT-2501 from 2018 competitive research projects call within the scope of PAIDI 2020—80% co-financed by the European Regional Development Fund (ERDF) from the Regional Ministry of Economic Transformation, Industry, Knowledge and Universities. Junta de Andalucía. Special thanks to the AECC (Spanish Association of Cancer Research) Founding Ref. GC16173720CARR for supporting this work. SMG was funded by a grant from the Fundación AECC. EMV-S and JMS-P are funded by postdoctoral fellowships from Junta de Andalucía (DOC_01655 and DOC_00512, respectively). The authors thank the donors and the HUVR-IBiS Biobank (Andalusian Public Health System Biobank and ISCIII-Red de Biobancos PT17/0015/0041) for the human specimens that were used in this study.

### Author contributions

SMG and AC conceived and designed this study. SMG and EMVS performed the experiments; JMSP analysed the NGS data; PEG collected the clinical data; and SMG and AC analysed and interpreted the data and drafted and edited the manuscript. All authors revised the manuscript.

## Notes

### Competing Interest Statement

The authors have declared no competing interest.

