## Supplementary Material for "Essential role of PLD2 in hypoxia-induced stemness and therapy resistance in ovarian tumors"

**Table S1. Reagents used in this work.**

| **Antibody** | **Dilution** | **Reference** |
|---|---|---|
| Anti-PLD2 (E1Y9G) | 1:1000 (Wb)/1:250(IF) | Cell Signaling ##13904 |
| Anti-HIF1A | 1:500 (Wb)/ 1:200 (IF) | CaymanChem#10006421 |
| Anti-SOX9 | 1:200 | Abcam # ab185230 |
| Anti-NOTCH1 | 1:250 | SantaCruz #sc-6014-r |
| Anti-Sox-2 (E-4) | 1:250 | SantaCruz # sc-365823 |
| Anti-SOX17 [EPR20684] | 1:250 | Abcam #ab224637 |
| mAb anti-α-tubulin | 1:5000 | Sigma 9026 |
| peroxidase-labeled rabbit anti-mouse | 1:10000 | Amersham |
| peroxidase-labeled goat anti-rabbit | 1:10000 | Abcam #6721 |
| **Probe** | | **Reference** |
| PLD2 | | ThermoFisher#Hs00160163 |
| LDHA | | ThermoFisher#Hs01378790 |
| VEGFA | | ThermoFisher#Hs00900055 |
| SOX2 | | ThermoFisher#Hs01053049 |
| NANOG | | ThermoFisher#Hs04260366 |
| CD44 | | ThermoFisher#Hs01075861 |
| EPCAM | | ThermoFisher#Hs00901885 |
| SOX9 | | ThermoFisher#Hs01001343 |
| NOTCH1 | | ThermoFisher#Hs01062014 |
| SNAI1 | | ThermoFisher#Hs00195591 |
| VIM | | ThermoFisher#Hs00958111 |
| CDH1 | | ThermoFisher#Hs01023894 |
| CDH2 | | ThermoFisher#Hs00983056 |
| ACTB | | ThermoFisher#Hs001060665 |

**Table S2: Patient Cohort characteristics**

| | Sensitive<br>N=10 (40%) | Resistant<br>N=15(60%) |
|---|---|---|
| **Age (years)**<br>• Mean (Rank) | 62,0 (34-70) | 51,0 (40-67) |
| **ECOG**<br>• 0<br>• 1<br>• 2 | 7 (77,8%)<br>1 (11,1%)<br>1 (11,1%) | 5 (38,5%)<br>6 (46,2%)<br>2 (15,4%) |
| **Stage (FIGO 2014)**<br>• IA<br>• IC<br>• IIB<br>• IIIB<br>• IIIC<br>• IVA<br>• IVB | 1 (11,1%)<br>1 (11,1%)<br>1 (11,1%)<br>1 (11,1%)<br>4 (44,4%)<br>1 (11,1%)<br>0 | 1 (7,7%)<br>1 (7,7%)<br>0<br>1 (7,7%)<br>8 (61,5%)<br>0<br>2 (15,4%) |
| **Ca 125 (U/ml)**<br>• **Diagnosis:** Median (Rank)<br>• **After treatment:** Median (Rank) | 194 (31,6-21957)<br>11 (6,5-1400) | 332 (38-3892)<br>76,1 (15,5-1862) |
| **Adjuvant Chemotherapy**<br>• No<br>• Yes | 6 (66,7%)<br>3 (33,3%) | 7 (53,8%)<br>6 (46,2%) |
| **Treatment**<br>• Carbo + Paclitaxel<br>• Carbo + Paclitaxel + beva<br>• Carbo monotherapy | 6 (68%)<br>3 (32%)<br>0 | 12 (92%)<br>0<br>1 (8%) |
| **Surgery**<br>• R0<br>• R1<br>• Biopsies<br>• No (incluyen pacientes con cirugía primaria) | 1 (11,1%)<br>2 (22,2%)<br>0<br>6 (66,7%) | 2 (15,4%)<br>2 (15,4%)<br>2 (15,4%)<br>7 (53,8%) |
| **Better response to QT adyuvant or 1st líne**<br>• RC<br>• RP | 6 (66,7%) | 4 (30,8%) |

| | | |
|---|---|---|
| • EE<br>• PE | 3 (33,3%)<br>0<br>0 | 2 (15,4%)<br>2 (15,4%)<br>5 (38,5%) |
| **Treatment after 1st line**<br>• Bevacizumab<br>• Others<br>• No | 3 (33,3%)<br>0<br>6 (66,7%) | 0<br>0<br>13 (100%) |
| **Progression disease after treatment**<br>• yes<br>• No | 6 (66,7%)<br>3 (33,3%) | 13 (100%)<br>0 |
| **Platinum free interval (moths)**<br>• Mean (Rank) | 19 (7-33) | 1 (0-5) |
| **More than 2 lines of treatment**<br>• yes<br>• No | 4 (44,4%)<br>5 (55,6%) | 4 (30,8%)<br>9 (69,12%) |
| **Status patient in last visit**<br>• Live without disease<br>• Live with disease<br>• Death (all due to disease progression) | 2 (22,2%)<br>4 (44,4%)<br>3 (33,3%) | 0<br>2 (15,4%)<br>11 (84,6%) |
| **Location primary tumor**<br>• Right ovary<br>• Left ovary<br>• Bilateral<br>• Peritoneal | 2 (22,2%)<br>2 (22,2%)<br>5 (55,6%)<br>0 | 2 (15,4%)<br>4 (30,8%)<br>5 (38,5%)<br>2 (15,4%) |
| **Differentiation**<br>• Moderately<br>• Poor<br>• nd | 1 (11,1%)<br>7 (77,8%)<br>1 (11,1%) | 1 (7,7%)<br>11 (84,6%)<br>1 (7,7%) |
| **Histology**<br>• Serous carcinoma<br>• Clear cell carcinoma<br>• Others | 7 (77,8%)<br>2 (22,2%)<br>0 | 8 (61,5%)<br>3 (23,1%)<br>2 (15,4%) |
| **Lymphovascular infiltration**<br>• No<br>• Yes<br>• nd | 1 (11,1%)<br>2 (22,2%)<br>6 (66,7%) | 1 (7,7%)<br>1 (7,7%)<br>11 (84,6%) |
| **BRCA Mutation** | | |

| | | |
|---|---|---|
| • No<br>• Yes<br>• nd | 4 (44,4%)<br>2 (22,2%)<br>3 (33,3%) | 6 (46,2%)<br>0<br>7 (53,8%) |

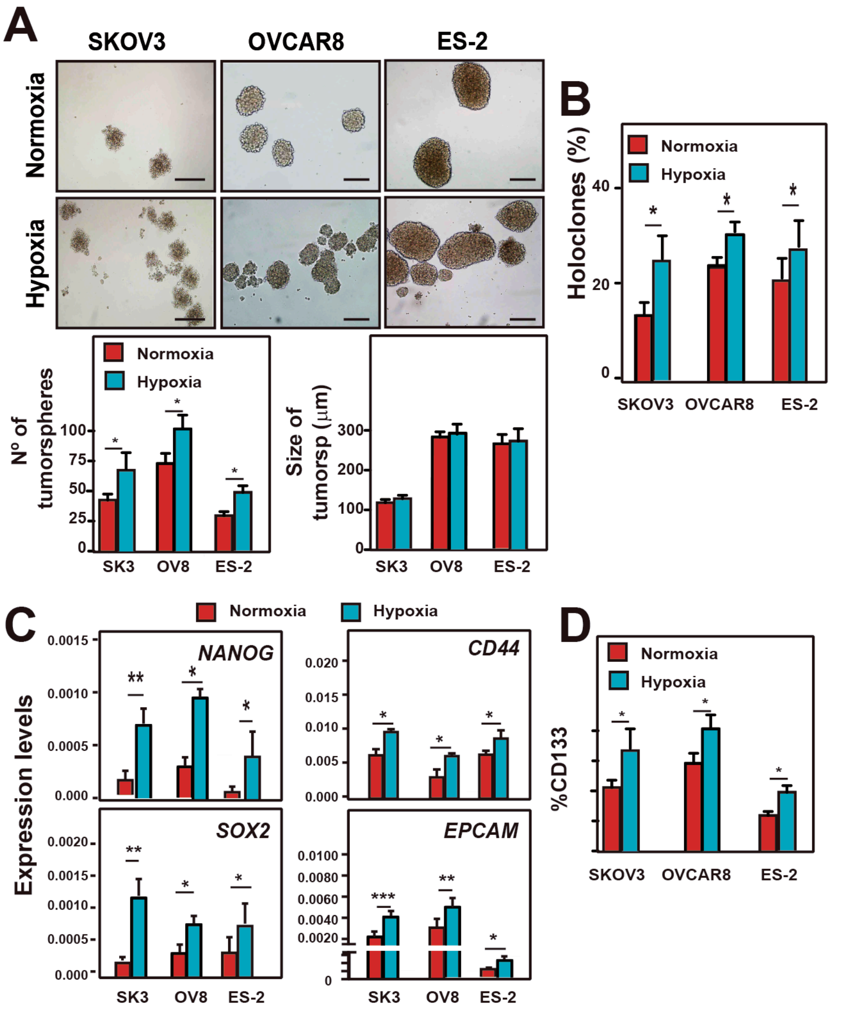


**Figure S1. Hypoxia induces CSCs in ovarian cancer cells. (A)** Top, Representative images of tumorspheres formed by SKOV3, OVCAR8 and ES-2 cells in normoxia or hypoxia. Bottom, quantification of the number and size of tumorspheres. Scale bars: 250 µm. **(B)** Percentage of holoclones formed by SKOV3, OVCAR8 and ES-2 cells in normoxia or hypoxia. At least 200 individual clones were analyzed. (**c**) Analysis of the expression of *NANOG*, *SOX2*, *CD44* and *EPCAM* stemness-associated genes by RT-qPCR in SKOV3, OVCAR8 and ES-2 cells in normoxia or hypoxia. The mRNA expression was calculated as $2^{-\Delta Ct}$ relative to the *ACTB* gene. The average and SD of three independent experiments are shown in all cases. A minimum of three independent experiments were performed and the data were compared using Student's t tests. Asterisks indicate statistical significance with respect to normoxia. $*p < 0.05$; $**p < 0.01$; $***p < 0.001$.

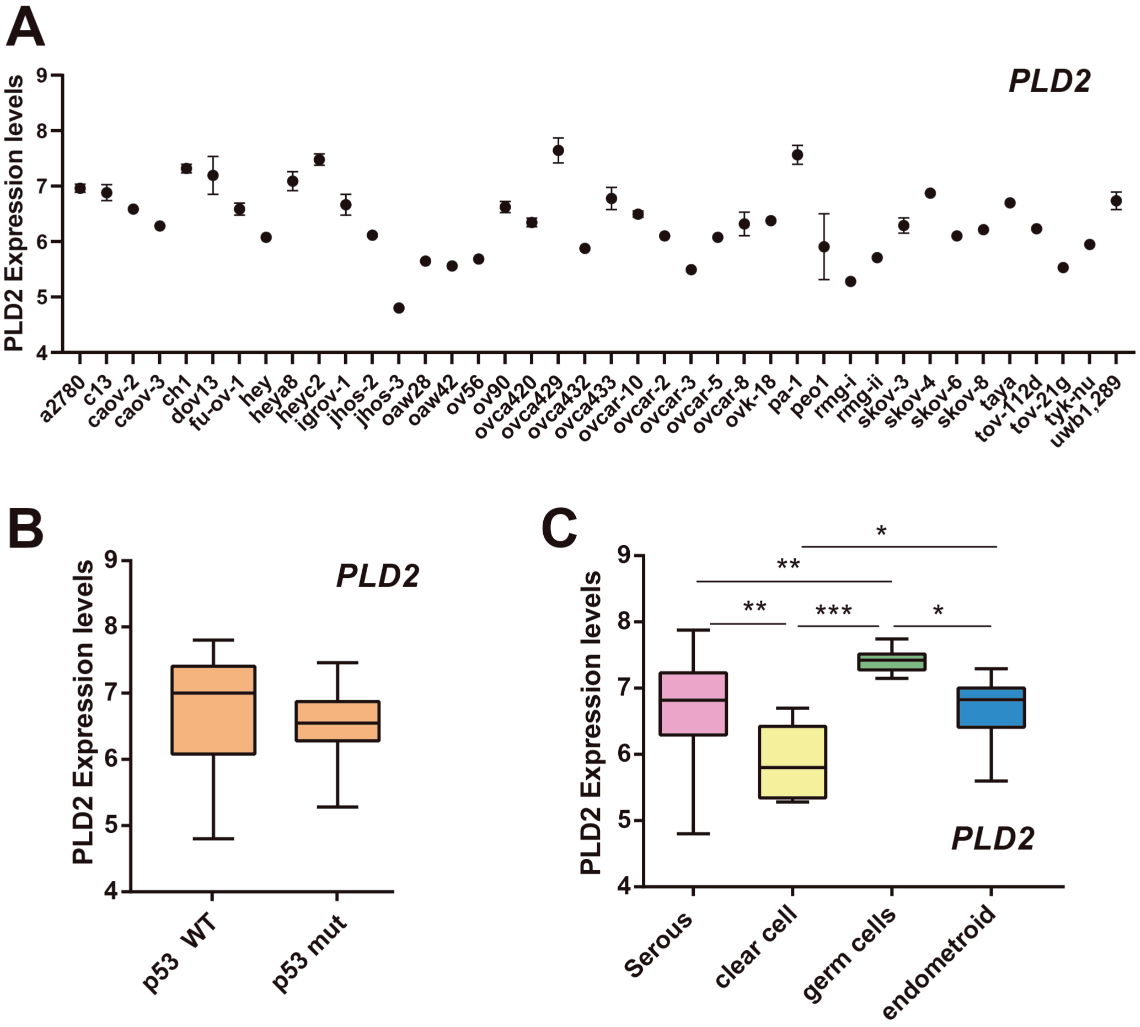


**Figure S2.** ***PLD2*** **expression in a panel of gynecological cancer cell lines. (A)** *PLD2* expression in a panel of gynecological cancer cell lines from GSE47856 (PMID 24858042). **(B)** *PLD2* expression in the cell lines from A splitted in two groups: p53 wild-type and mutant cell lines. (**C**) *PLD2* expression in the cell lines from A depending on their histotype. Asterisks indicate statistical significance according to Student's t tests. $^*p < 0.05$; $^{**}p < 0.01$; $^{***}p < 0.001$.

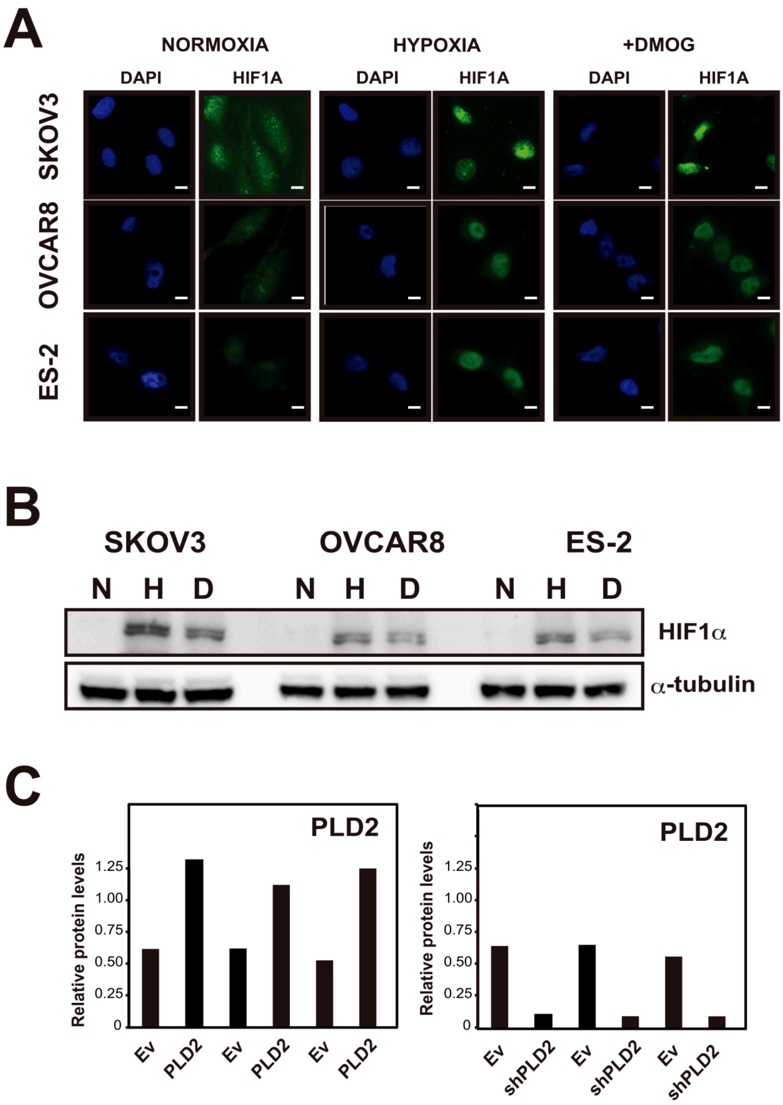


**Figure S3. (A)** Representative images of HIF-1α protein levels by immunofluorescence in SKOV3, OVCAR8 and ES-2 cells under normoxia and hypoxia or in the presence of the HIF-hydroxylase inhibitor DMOG. **(B)** Western blot showing HIF-1α, and alpha-tubulin protein levels in SKOV3, OVCAR8 and ES-2 ovarian cancer cells under normoxia and hypoxia or in the presence of the HIF-hydroxylase inhibitor DMOG. **(C)** Relative protein quantification of PLD2 normalized to alpha-tubulin from the western blots in Figure 2B. A minimum of three independent experiments were performed and the data were compared using Student's t tests. *$p < 0.05$; **$p < 0.01$; ***$p < 0.001$.

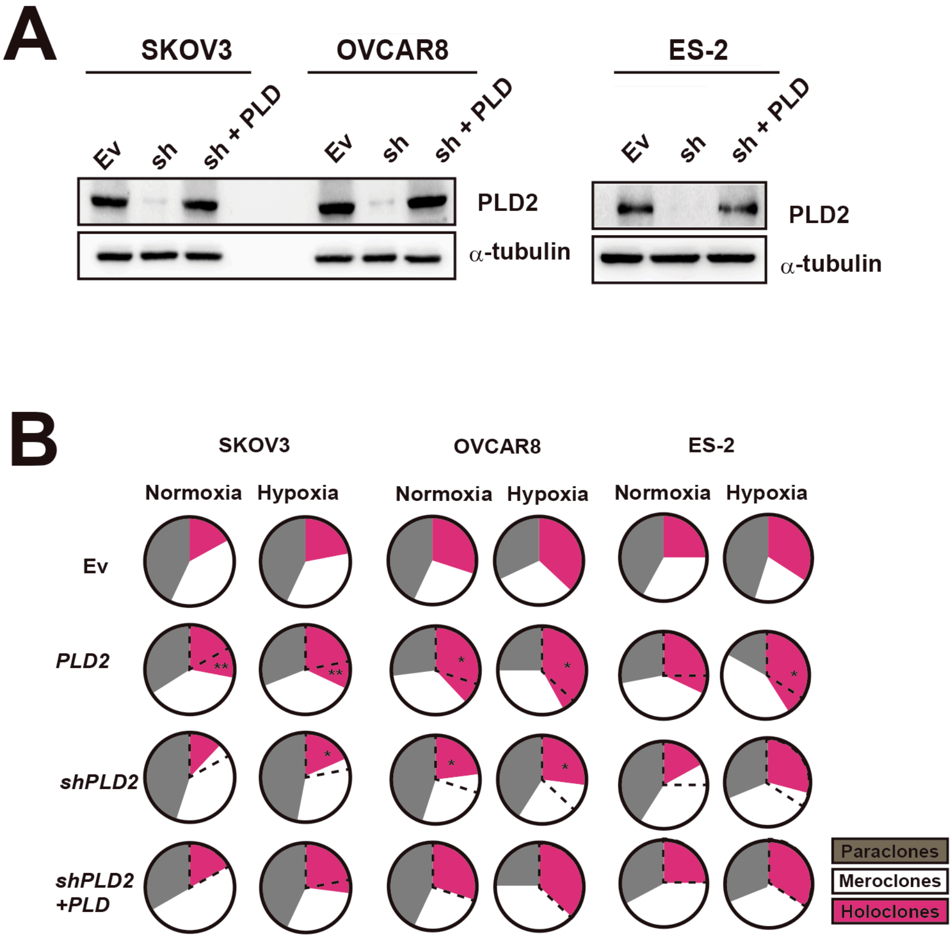


**Figure S4.** (**A**) Western blot showing PLD2 and alpha-tubulin protein levels in SKOV3, OVCAR8 and ES-2 OC cells carrying Ev, a plasmid expressing *shPLD2* or plasmids expressing *shPLD2* and *PLD2*. (**B**) Percentage of paraclones, meroclones and holoclones formed by SKOV3, OVCAR8 and ES-2 cells carrying Ev or plasmids expressing *PLD2*, *shPLD2* or both in hypoxia or normoxia. At least 200 individual clones were analyzed. The average of three independent experiments is shown. A dotted line represents the percentage of holoclones in Ev carrying cells as a reference. Data were compared using Student's t tests. Asterisks indicate statistical significance with respect to Ev carrying cells. *p < 0.05.

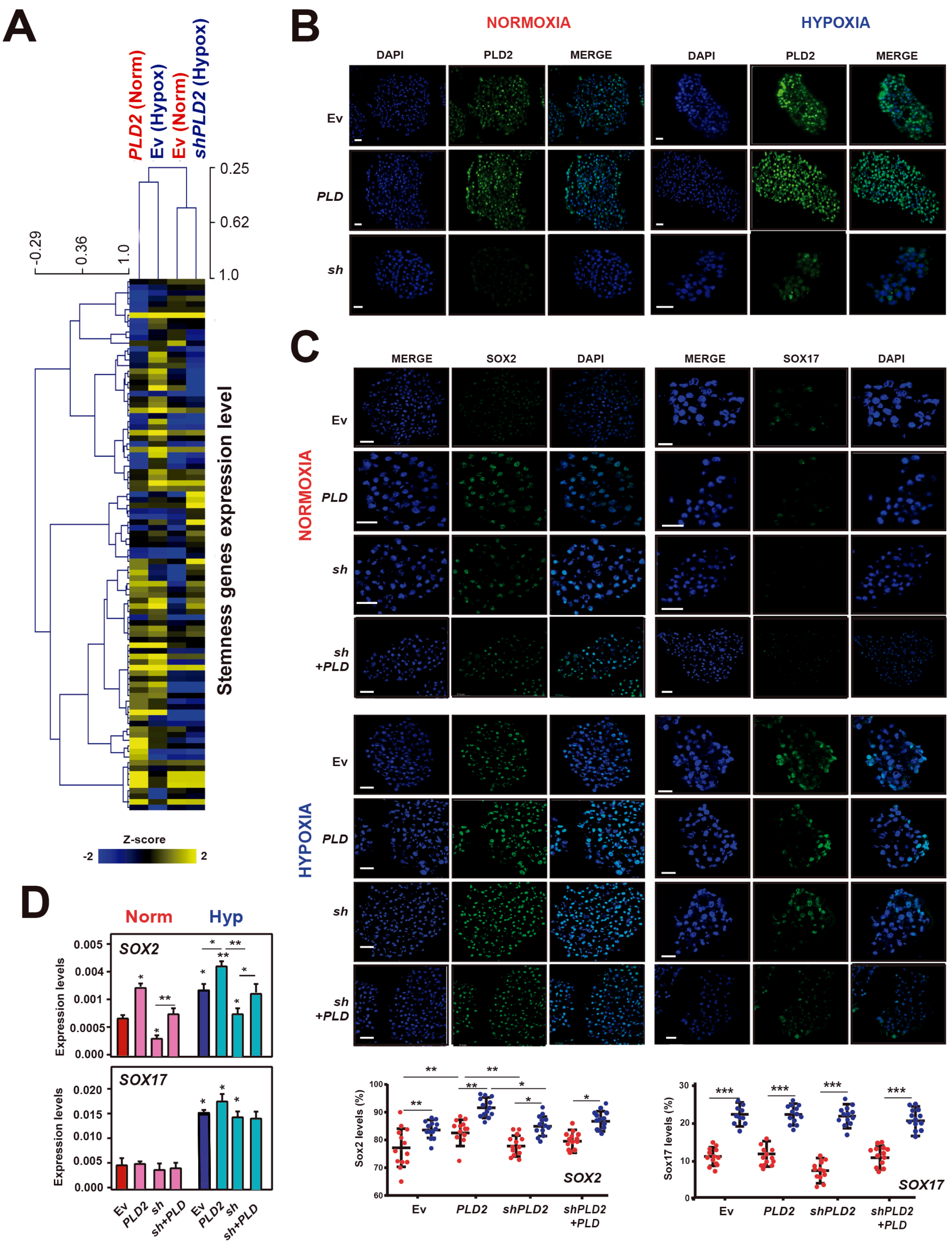


**Figure S5. (A)** Heatmaps showing the z-scores of stemness genes expression levels in SKOV3 cells carrying EV or plasmid overexpressing *PLD2* under normoxia and carrying EV or plasmid expressing *shPLD2* under hypoxia. Hierarchical clustering of the samples is shown. **(B)** Determination of PLD2 protein levels in tumorspheres formed by OC cell lines carrying Ev or plasmids expressing *PLD2* or *shPLD2*. Scale bars: 100 µm. **(C)** Top, determination of Sox2 and Sox17 protein levels in tumorspheres formed by OC cell lines carrying Ev or plasmids expressing *PLD2*, *shPLD2* or both. Scale bars: 100 µm. Bottom, representative quantification of the percentage of cells with Sox2 and Sox17 expression in tumorspheres formed by OC cell lines carrying Ev or plasmids expressing *PLD2*, *shPLD2* or both. **(D)** Expression levels by RT-qPCR of *SOX2* and *SOX17* stemness associated genes in tumorspheres formed by OC cell lines carrying Ev or plasmids expressing *PLD2*, *shPLD2* or both. The mRNA expression was calculated as $2^{-\Delta Ct}$ relative to the *ACTB* gene. A minimum of three independent experiments were performed and the data were analyzed using Student's t-test. *, $P < 0.05$; **, $P < 0.01$; ***, $P < 0.001$.

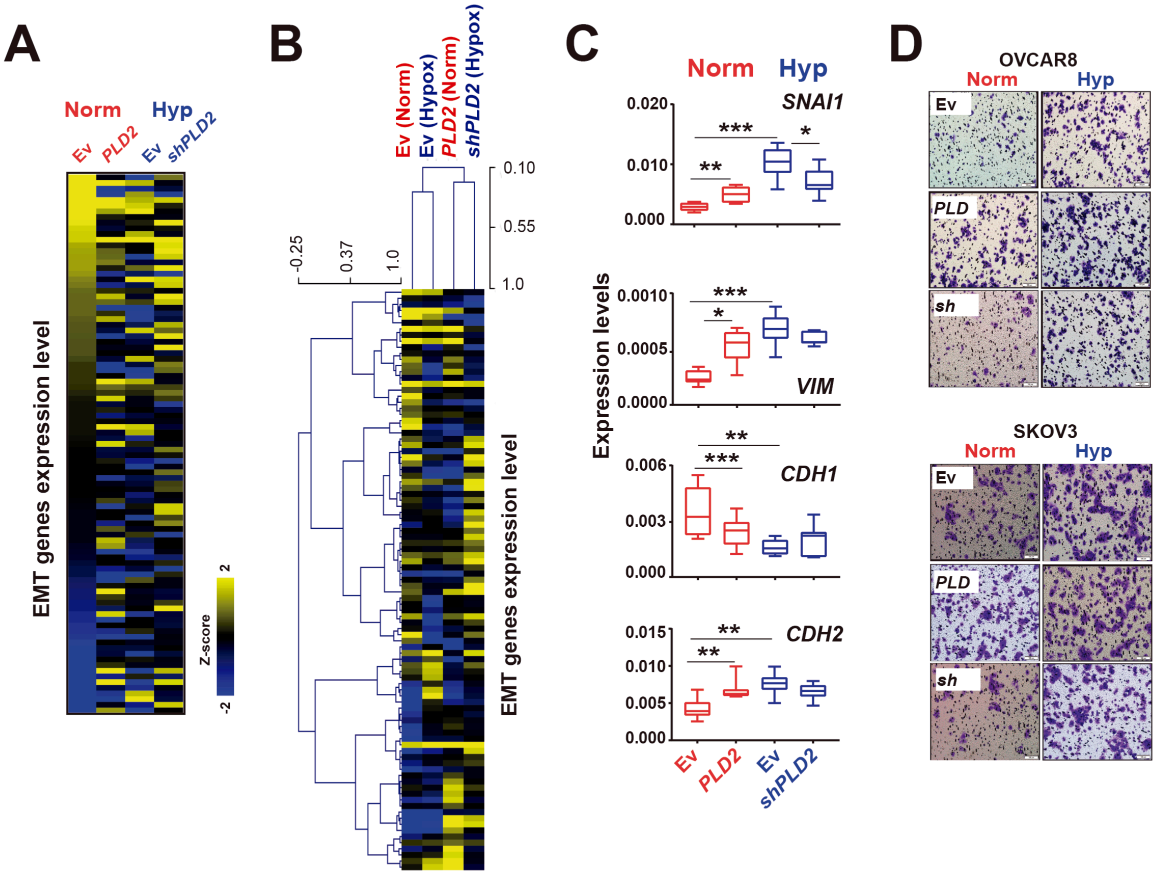


**Figure S6. (A)** Heatmaps showing the expression z-scores of epithelial-to-mesenchymal transition associated genes obtained from TaqMan Arrays. Genes are sorted according to decreasing z-scores in the Ev-carrying cells under normoxia. **(B)** Heatmaps showing the z-scores of EMT genes expression levels in SKOV3 cells carrying EV or plasmid overexpressing *PLD2* under normoxia conditions and carrying EV or plasmid expressing *shPLD2* under hypoxia condition. Hierarchical clustering of the samples is shown. **(C)** Expression levels of *SNAI1, VIM, CDH1* and *CDH2* EMT-associated genes in cells carrying Ev or plasmids expressing *PLD2* under normoxia or *shPLD2* under hypoxia conditions. **(D)** Representative images of the Boyden chamber migration assays in SKOV3 and OVCAR8 cells carrying Ev or plasmids expressing *PLD2* or *shPLD2* under normoxic or hypoxic conditions from Figure 3F. A minimum of three independent experiments were performed and the data were analyzed using Student's *t*-test. Asterisks indicate statistical significance with respect to Ev carrying cells in normoxia, unless indicated by horizontal lines. *, $P < 0.05$; **, $P < 0.01$; ***, $P < 0.001$.

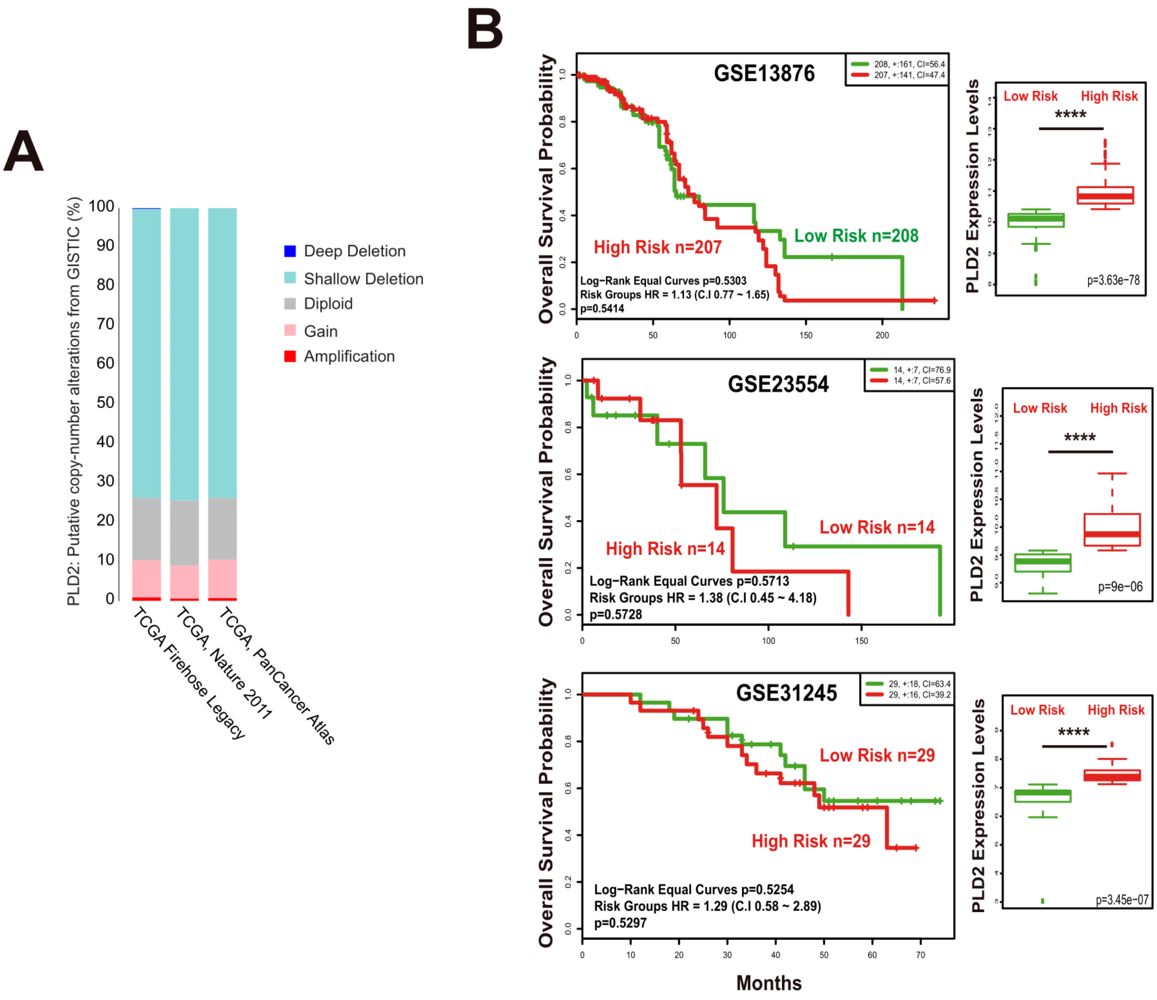


**Figure S7. (A)** Quantification of copy number alterations in OC patients from the TCGA database in the studies PanCancer Atlas, TCGA Nature 2011, and TCGA Firehouse legacy. Data were obtained from cBioportal. **(B)** Kaplan-Meier plots showing overall survival of patients with high (red) or low (green) *PLD2* expression levels in three databases with survival data (GSE13876, GSE23554 and GSE31245). Data were analyzed with the log-rank test, and the associated P-values are shown in the graphs. Expression levels are shown as $\log_2$ transformed values from the R2 database. Data were analyzed using Student's *t*-test. *, P < 0.05; **, P < 0.01; ***, P < 0.001.

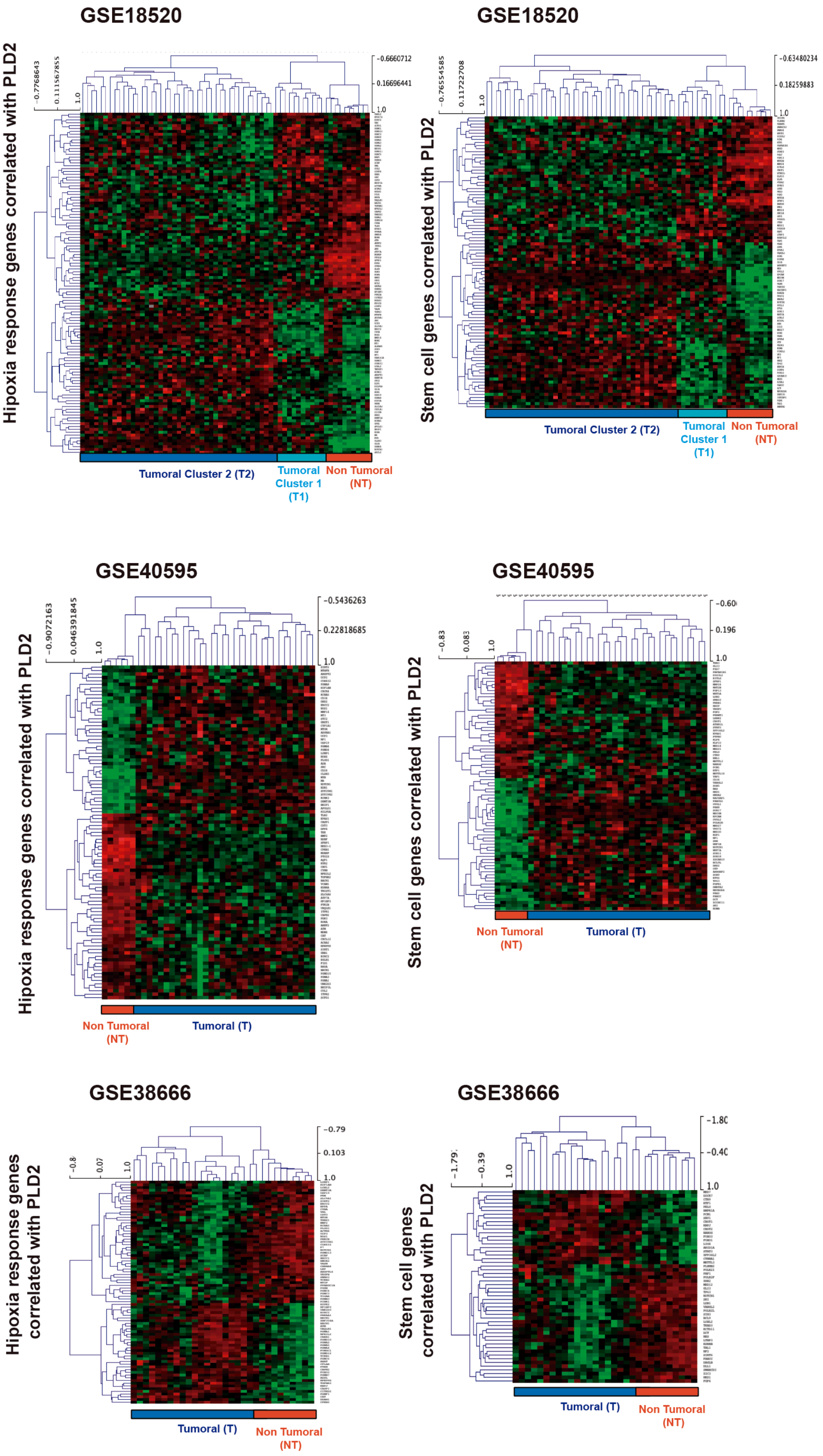


**Figure S8.** Heatmaps showing the expression z-scores of stemness-associated genes or hypoxia-response genes whose expression correlated with *PLD2* in GSE18520, GSE38666 and GSE40595 OC patient databases.

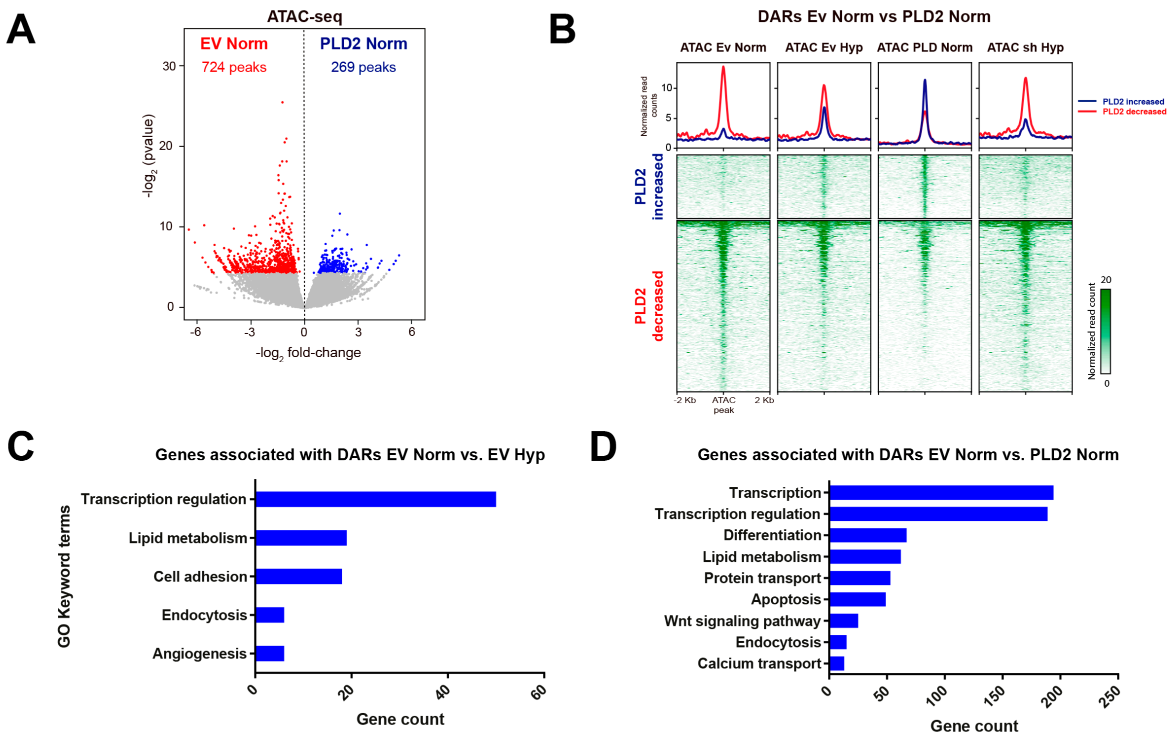


**Figure S9. (A)** Volcano plot showing differential analyses of chromatin accessibility between SKOV3 cells carrying Ev and plasmid overexpressing *PLD2* in normoxia. **(B)** Heatmaps and average profiles plotting normalized ATAC-seq signal in SKOV3 cells carrying Ev and overexpressing *PLD2* for the differentially accessible regions (DARs) in (A). **(C)** Gene Ontology term enrichment analyses of biological processes for the genes associated with DARs in SKOV3 cells carrying EV in normoxia versus hypoxia. **(D)** Gene Ontology term enrichment analyses of biological processes for the genes associated with DARs in SKOV3 cells carrying Ev versus overexpressing *PLD2* in normoxia.

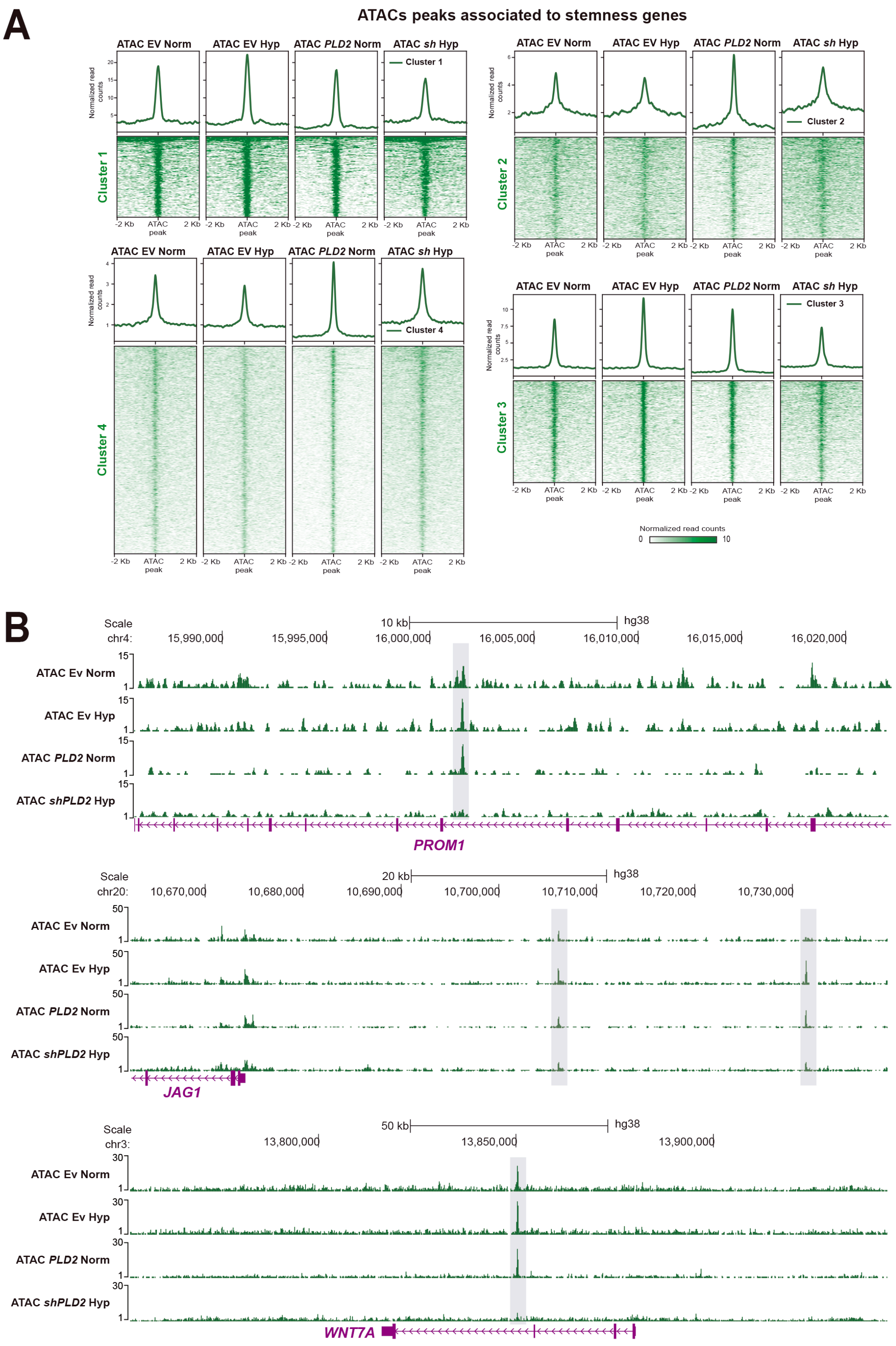


**Figure S10. (A)** Heatmaps plotting normalized ATACs-seq signal at peaks associated with stemness genes in SKOV3 cells carrying Ev or plasmid expressing *PLD2* in normoxia and carrying Ev or plasmid expressing *shPLD2* in hypoxia, for the differentially accessible regions (DARs) clustered using *k*-means method in 4 clusters. **(B)** Tracks with ATAC-seq in SKOV3 cells carrying Ev or expressing *PLD2* in normoxia and carrying Ev or *shPLD2* in hypoxia, at the *PROM1, JAG1* and *WNT7A* loci.

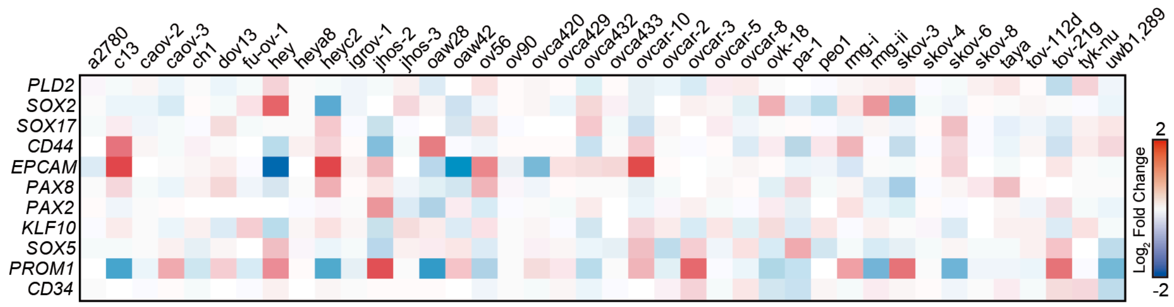


**Figure S11. (A)** Heatmap showing the $\log_2$ fold change of expression of cisplatin treated vs. untreated cells for *PLD2* and several stemness-associated genes in a panel of gynecological cancer cell lines from GSE47856 (PMID 24858042).
